# Comparing dormancy in two distantly related tunicates reveals morphological, molecular, and ecological convergences and repeated co-option

**DOI:** 10.1101/2022.02.09.477513

**Authors:** Laurel S. Hiebert, Marta Scelzo, Alexandre Alié, Anthony De Tomaso, Federico Brown, Stefano Tiozzo

## Abstract

Many asexually-propagating marine invertebrates can survive extreme environmental conditions by developing dormant structures, i.e., morphologically simplified bodies that retain the capacity to completely regenerate a functional adult when conditions return to normal. Here, we examine the environmental, morphological, and molecular characteristics of dormancy in two distantly related clonal tunicate species: *Polyandrocarpa zorritensis* and *Clavelina lepadiformis*. In both species, we report that the dormant structures are able to withstand harsher temperature and salinity conditions compared to the adult, and are the dominant forms these species employ to survive the colder winter months. By finely controlling the entry and exit of dormancy in laboratory-reared individuals, we were able to select and characterize the morphology of dormant structures associated with their transcriptome dynamics. In both species, we identified putative stem and nutritive cells in structures that resemble the earliest stages of asexual propagation. By characterizing gene expression during dormancy and regeneration into the adult body plan (i.e., germination), we observed that genes which control dormancy and environmental sensing in other metazoans, notably HIF-α and insulin signaling genes, are also expressed in tunicate dormancy. Germination-related genes in these two species, such as the retinoic acid pathway, are also found in other unrelated clonal tunicates during asexual development. These results are suggestive of repeated exaptation of conserved eco-physiological and regeneration programs for the origin of novel dormancy-germination processes across distantly related animal taxa.

## Introduction

To survive unfavorable environmental conditions, many animal species have evolved a remarkable strategy that allows them to reduce their metabolic rate and enter into a resting state; a process called dormancy (reviewed by Hand 1991; Cáceres 1997; Wilsterman et al 2021). Molecular and physiological mechanisms underlying dormancy across a variety of contexts in metazoans point to the existence of recurring processes (Bertolani et al 2019; Guidetti et al 2011; Hahn and Denlinger, 2011; Ragland et al 2010; Wang et al 2009; Dias et al 2021). For instance, the initial phases of dormancy involve the arrest of any ongoing development or cell proliferation and the onset of hypometabolism (Storey and Storey 2010). Stress response gene expression increases, enhancing stress tolerance and stabilizing cellular components. Metabolism shifts from carbohydrate to lipid combustion, and anabolism (such DNA replication, transcription and translation) is radically attenuated (Hand et al 2016; Dias et al 2021). During the maintenance phase of dormancy molecular chaperones, antioxidants, proteins responsible for DNA and chromatin stabilization or repair are produced, conferring environmental stress resistance (Hand et al 2016; Dias et al 2021).

Among invertebrates, dormancy is found in 67 classes from 29 free-living phyla, and, depending on the species, can occur at different phases of their life cycle (Cáceres, 1997). Many of the species that utilize dormancy are also clonal, meaning they grow via repeated rounds of asexual reproduction during which entire bodies are regenerated. In these clonal species, the dormant form has a morphologically simpler body plan that can survive in conditions that the feeding adults cannot. The dormant structures in clonal species include: podocysts in scyphozoan cnidarians (Ikeda, Ohtsu, and Uye 2011), statoblasts in freshwater bryozoans (Bushnell and Rao 1974), hibernaculae in kamptozoans (Hyman 1940; Mukai and Makioka 1978), gemmules in sponges (Hand 1991), and “survival buds’’ or “winter buds” (among other names) in tunicates (Nakauchi 1982; Hyams et al 2017).

In species that couple dormancy with asexual propagation, the dormant structures themselves are generally morphologically simple, but contain progenitor and nutrient storage cells that can initiate and support germination. For example, bryozoan statoblasts can survive desiccation, freezing, or even pass through the digestive tracts of aquatic vertebrates unharmed (Brown 1933; Bushnell and Rao 1974). The statoblast consists of a germinal mass inside two sclerotized valves containing an outer layer of epithelial cells and an inner mass of yolk (Mukai 1982). During germination, the epithelial cells undergo a thickening followed by invagination, and subsequent development follows that of typical bryozoan budding (Wood 2015). In some Demospongiae species, gemmules develop inside the sponge tissues. They are made up of a collagenous coat with siliceous spicules. Inside the coat are a few thousand pluripotent cells, each packed with nutrients in the form of yolk inclusions. During germination, cells inside migrate out of the coat, proliferate and differentiate to become juvenile sponges (Simpson 1984). Only when the sponge dies do the gemmules have the capacity to germinate (Simpson 1984). However, gemmules can remain dormant under certain low temperatures when separated from the rest of the sponge (Simpson 1984). The stimulus responsible for germination appears to differ greatly across clonal dormancy-capable species, but typically germination occurs after separation from the adult under the proper environmental conditions.

To better understand the chain of events that span from the environmental triggering signals to the morphological and molecular changes that occur during the entrance and exit of a dormant state, we focused on the chordate subphylum Tunicata. In tunicates, dormancy has been reported exclusively in clonal species, i.e., species that undergo asexual reproduction by different types of budding (Alié et al. 2020) (Figure 1). In clonal tunicates, budding is driven either by circulatory stem cells and/or by the ability of some epithelia to transdifferentiate (Brown and Swalla 2012; Kawamura and Fujiwara 1995; Kassmer et al 2019; Freeman 1964; Kürn et al 2011; Sköld et al 2009). While the mechanisms underlying budding differ across species and often involve nonhomologous tissues and cells (Alié et al 2020, Brown and Swalla 2012), their development generally converges at an early double-layered vesicle stage, reminiscent of a blastula, that eventually forms into a new feeding zooid (Alié et al 2020, Tiozzo et al 2008). In the colonial species for which a dormant stage has been documented, dormancy occurs by arresting the budding process (Mukai et al 1983; Nakauchi 1982; Hyams et al 2017, Huxley 1921; Mukai et al 1983; Huxley 1926).

**Figure 1:**
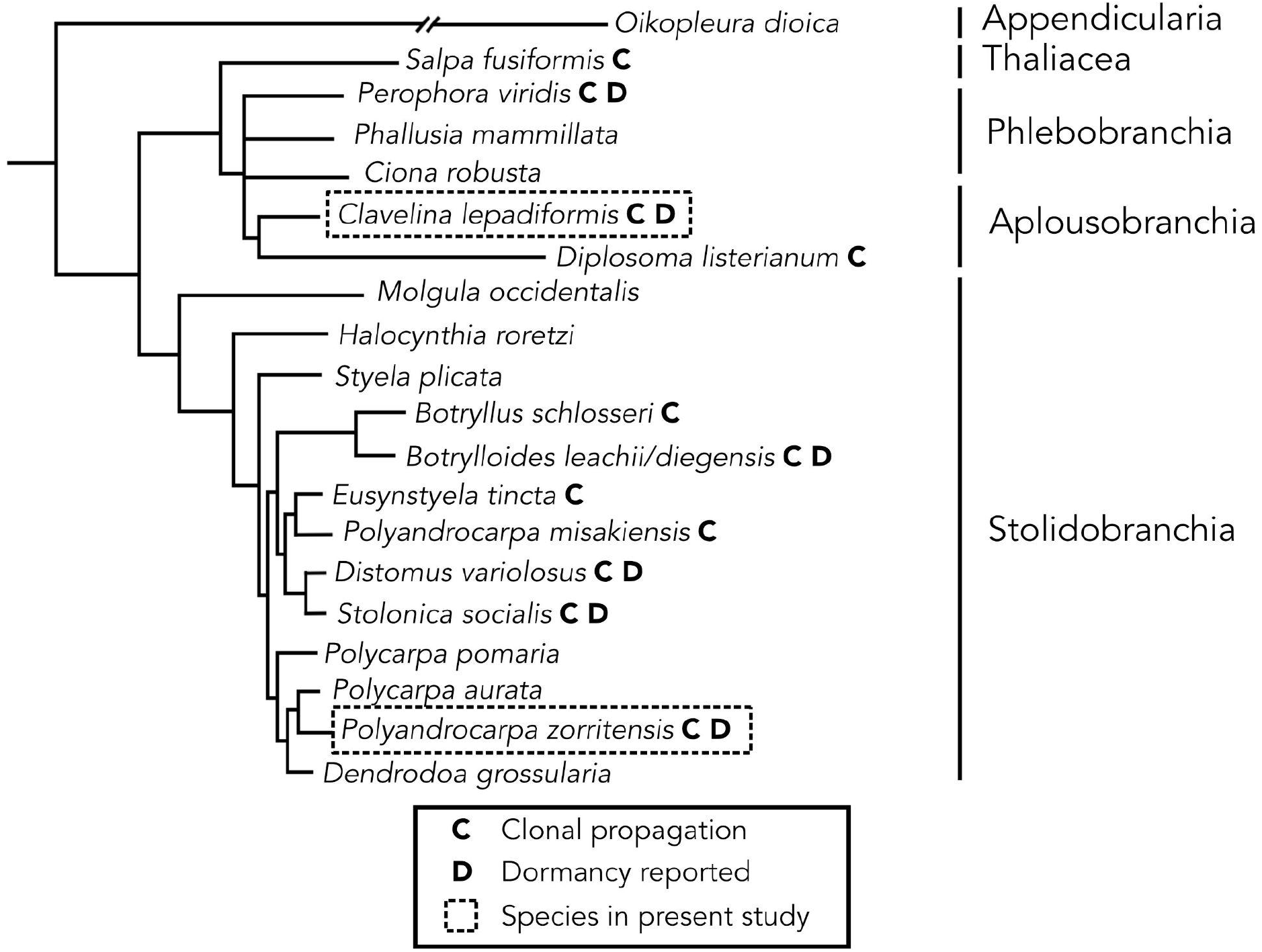
Consensus phylogeny of selected tunicate species indicating correlation between asexual cloning and dormancy. Orders are shown on the right. Species with clonal propagation are shown with “C” next to name. Species with capacity for dormancy are shown with a “D” next to name. The species in this study are boxed in a dashed line. Branch lengths and relationships are approximated from Alié et al. (2018) (for Stolidobranchia), Delsuc et al. (2018) (for Appendicularia, Thaliacea, Aplousobranchia, and Phlebobranchia except for *Perophora*), Tsagkogeorga et al. (2009) (for position of *Perophora*).

In this study we chose two relatively distant species belonging to two tunicate orders: *Polyandrocarpa zorritensis* (Stolidobranchia) and *Clavelina lepadiformis* (Aplousobranchia) (Figure 2-4), we described their seasonal behaviour in natural and laboratory conditions and analyzed their entrance and exit from the state of dormancy via anatomical and molecular characterizations. During *P. zorritensis* and *C. lepadiformis* asexual life-cycles (Figure 2), epidermal projections of the zooids, called stolons, begin to ramify forming structures called “budding nests” (as in *P. zorritensis*) or to lobulate forming structures called “budding chambers” (as in *C. lepadiformis*), here for simplicity both referred as “pre-buds.” Pre-buds can either give rise directly to buds and eventually develop into adult zooids, or, they can turn into dormant structures, orange spherical-like shapes known as “spherules” in *P. zorritensis* (Figure 2A; Scelzo et al., 2019), and white opaque dilations known as “winter buds” in *C. lepadiformis* (Figure 2B; Berrill, 1951). Winter buds have been previously reported in the Atlantic and Mediterranean in *C. lepadiformis* (Giard and Caullery, 1896; Orton 1921; Della Valle 1915: Huxley 1926). In this study we document for the first time seasonal variations of a *P. zorritensis* population in a temperate latitude and report the effect of temperature and salinity on the induction and release of dormancy in both species. We characterize and compare the morphological cellular structure and the transcriptomic profiles of dormant forms between these two species, and in order to better understand the molecular mechanisms involved in entry, maintenance, and release of dormancy, we present here the first results of a differential gene expression analysis of a number of stages of the life cycle of both species. Gene expression changes during dormancy in tunicates has not been previously studied.

**Figure 2:**
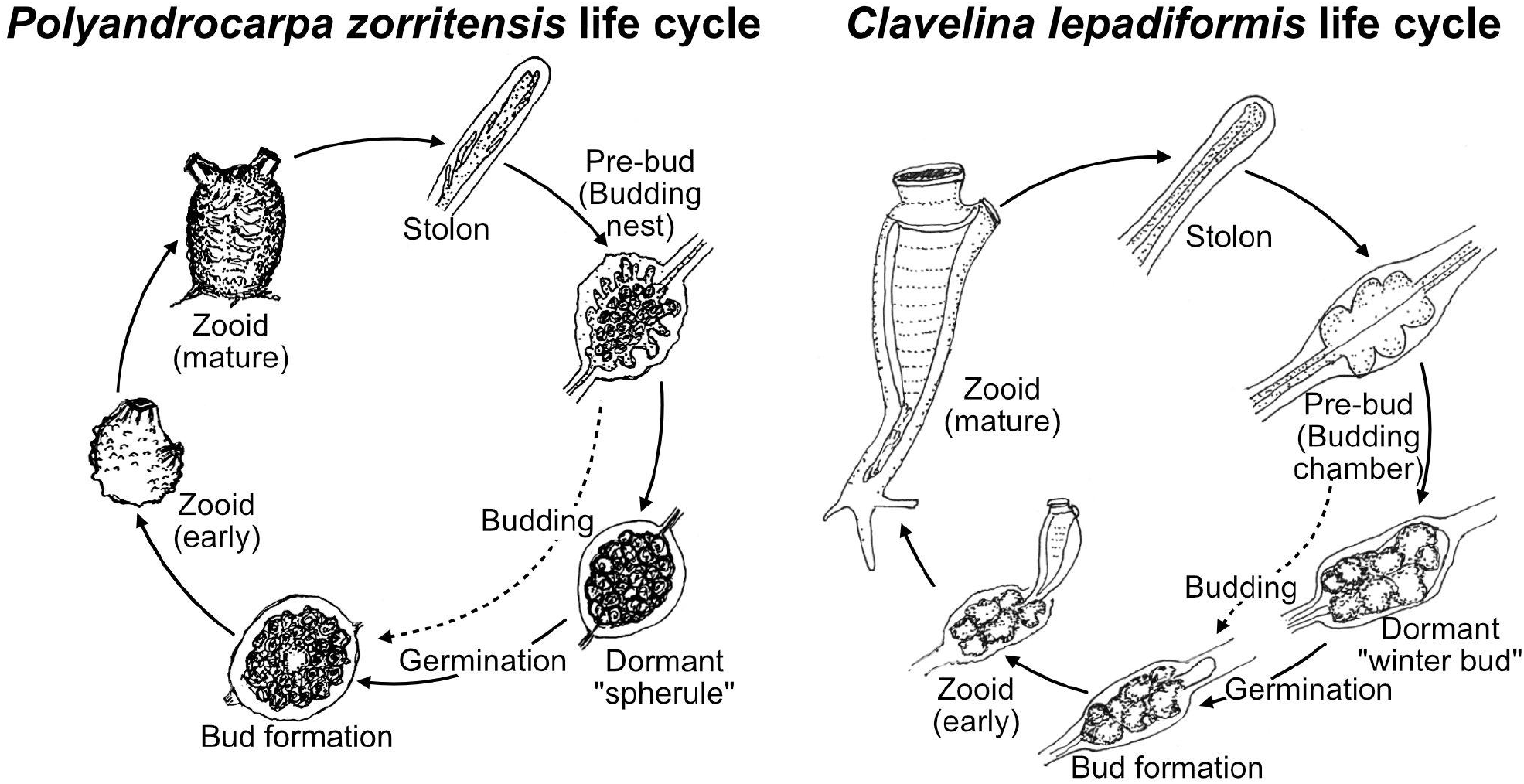
Asexual life cycles of *Polyandrocarpa zorritensis* and *Clavelina lepadiformis*. At the base of mature zooids of both *P. zorritensis* and *C. lepadiformis*, extensions of the blood vessels encased in tunic, called stolons, protrude along the substrate. Along the stolons, ramifications of the internal blood vessels develop that we call “pre-buds” here (and are also known as “budding nests” in *P. zorritensis* and “budding chambers” in *C. lepadiformis*). The pre-buds swell and the surrounding tunic thickens to form structures called “spherules” for *P. zorritensis* or may accumulate opaque mass inside the vessel to become what has been called “winter buds” in *C. lepadiformis*. These dormant forms go on to germinate into a new zooid, but they have the capacity to withstand harsh conditions before germinating. Note that the pre-bud can also initiate the budding process without passing through a dormant state, as indicated by the dotted line in each life cycle.

**Figure 3:**
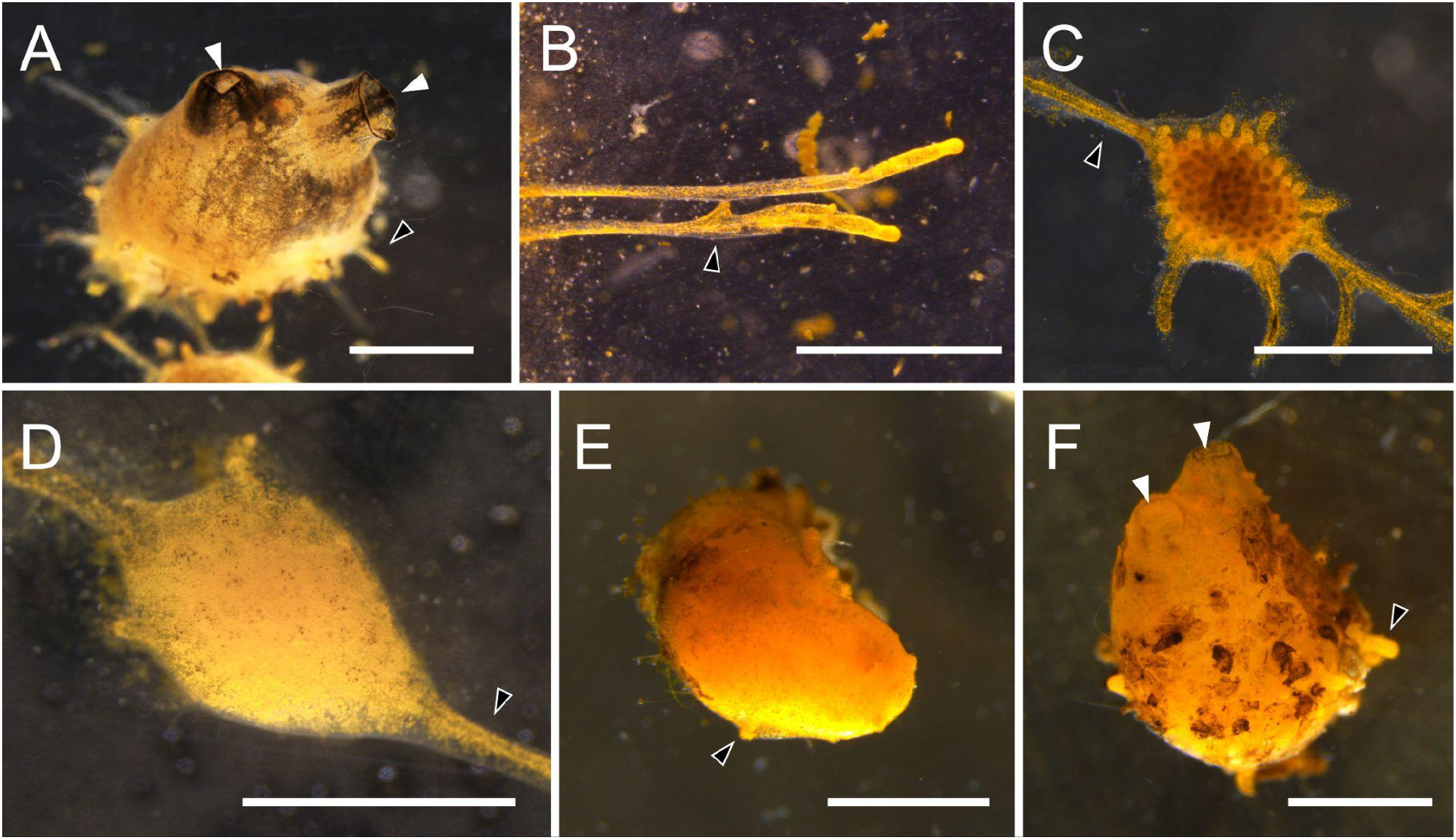
Photomicrographs of *Polyandrocarpa zorritensis* asexual stages. (A) Zooid. Siphons (white arrowheads) open for feeding. Stolons (black arrowhead marks one as an example) help attach the zooid to the substrate, and have begun to grow laterally. (B) Stolons attached firmly on a glass slide. Lower stolon starting to ramify (black arrowhead) where new pre-bud will likely develop. (C) Pre-bud still attached to “parental” zooid by stolon (black arrowhead), thus no bud tissues have begun to develop. Ramifications of vasculature seen within the structure through the tunic. (D) Early spherule, as indicated by thickened tunic, which obscures the vascular ramifications inside. This spherule remains attached to the zooid by the stolon, and thus budding has not yet initiated. (E) Detached spherule undergoes germination, as stolonal outgrowths form (black arrowhead). (F) Young zooid that arose from a hatched spherule. Siphons are open (white arrowheads), suggesting that filter feeding has been initiated. Stolon (black arrowhead) begins to grow outward. Scale bars are all 1mm.

**Figure 4:**
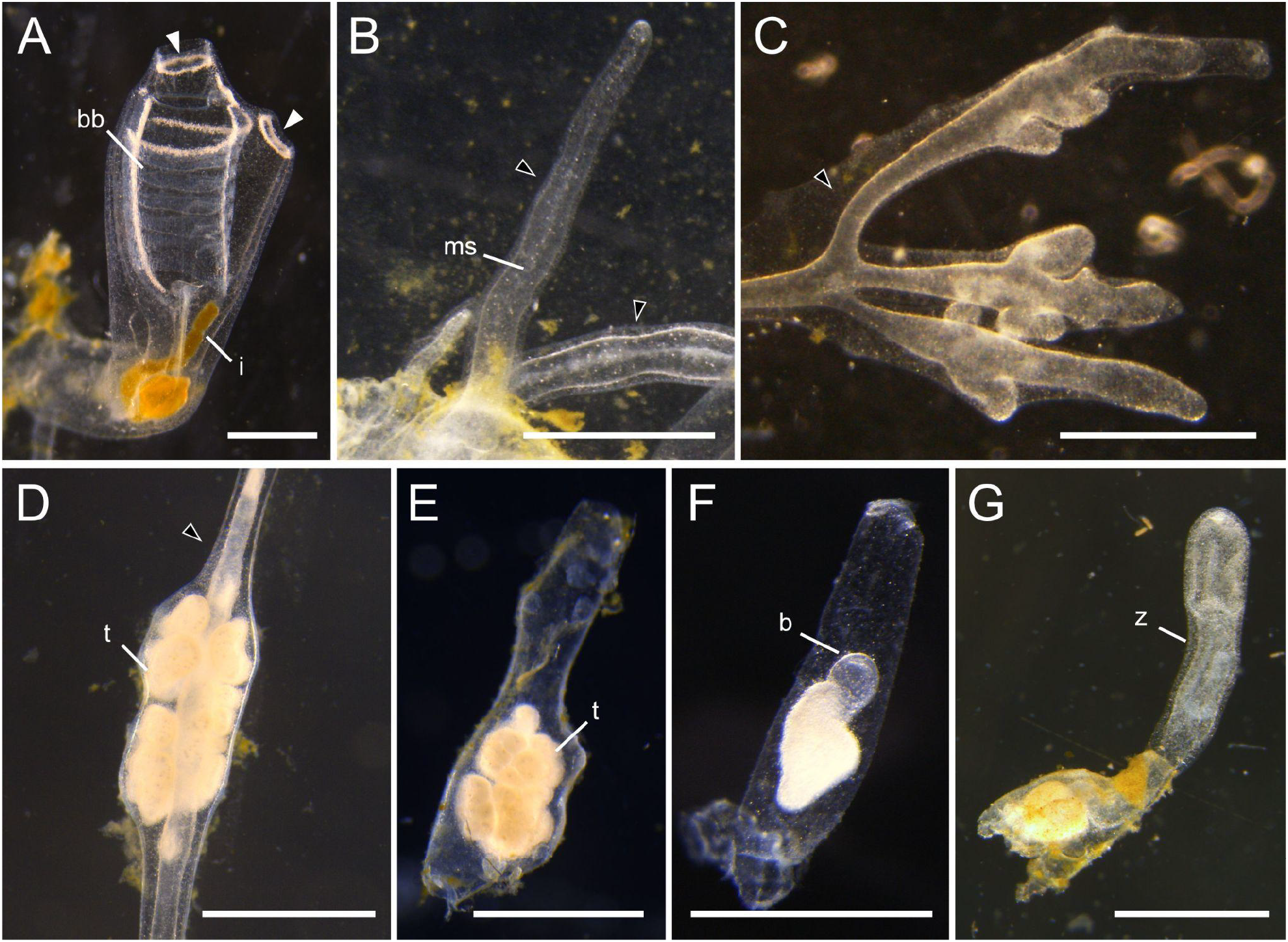
Photomicrographs of *Clavelina lepadiformis* asexual stages. (A) View of a feeding zooid with siphons (white arrowheads), branchial basket (“bb”), and intestine (“i”). (B) Stolons (marked with black arrowheads) protruding from zooid allow for attachment to substrate and for production of pre-buds. Mesenchymal septum (“ms”) indicated within stolon. (C) Stolons (marked with black arrowheads) undergo branching morphogenesis near the tips and begin to form the pre-buds (also called budding chambers in this species). (D) Early winter bud in a stolon still attached to the zooid. It has accumulated mass of trophocytes (“t”) inside as indicated by opaque white material. Stolon marked with white arrowhead. (E) Detached winter bud with a large mass of trophocytes (“t”). (F) Winter bud undergoing early steps of germination. The germinating bud (b) seen as a small transparent bump above the trophocyte mass. (G) Late germination stage showing young zooid (“z”) that arose from winter bud. Siphons are not yet visible. Remnants of winter bud remain and continue to supply nutrition to zooid until it begins to feed on its own or until it runs out of the supplied nutritive material. Scale bars are all 1mm.

## Results

### 1. P. zorritensis *spherule and zooid turnover conforms with seasonal changes*

To evaluate the presence of dormant versus actively growing stages in their natural habitat, a population of *P. zorritensis* located in La Spezia (Italy) was monitored over two years (from August 2017 to May 2019) with two to three observations per season (Figure 5). In summer, fall, and winter the colonies formed dense clusters of zooids, and dormant spherules were connected to the colony by stolons on the bottom sides of these clusters (Figure 5B). During two springs (March 2018 and May 2019) no zooids were observed and the colony consisted solely of isolated spherules (as in Figure 3E and 5C), attached to the boat mooring lines, and covered by epibionts. Spherules collected over the year (2018-2019) showed differences in average diameter (Figure 5C-D). The average spherule size was significantly larger during fall than in winter, and significantly larger in winter than spring (RM-ANOVA, p<0.03; Figure 5C-D).

**Figure 5:**
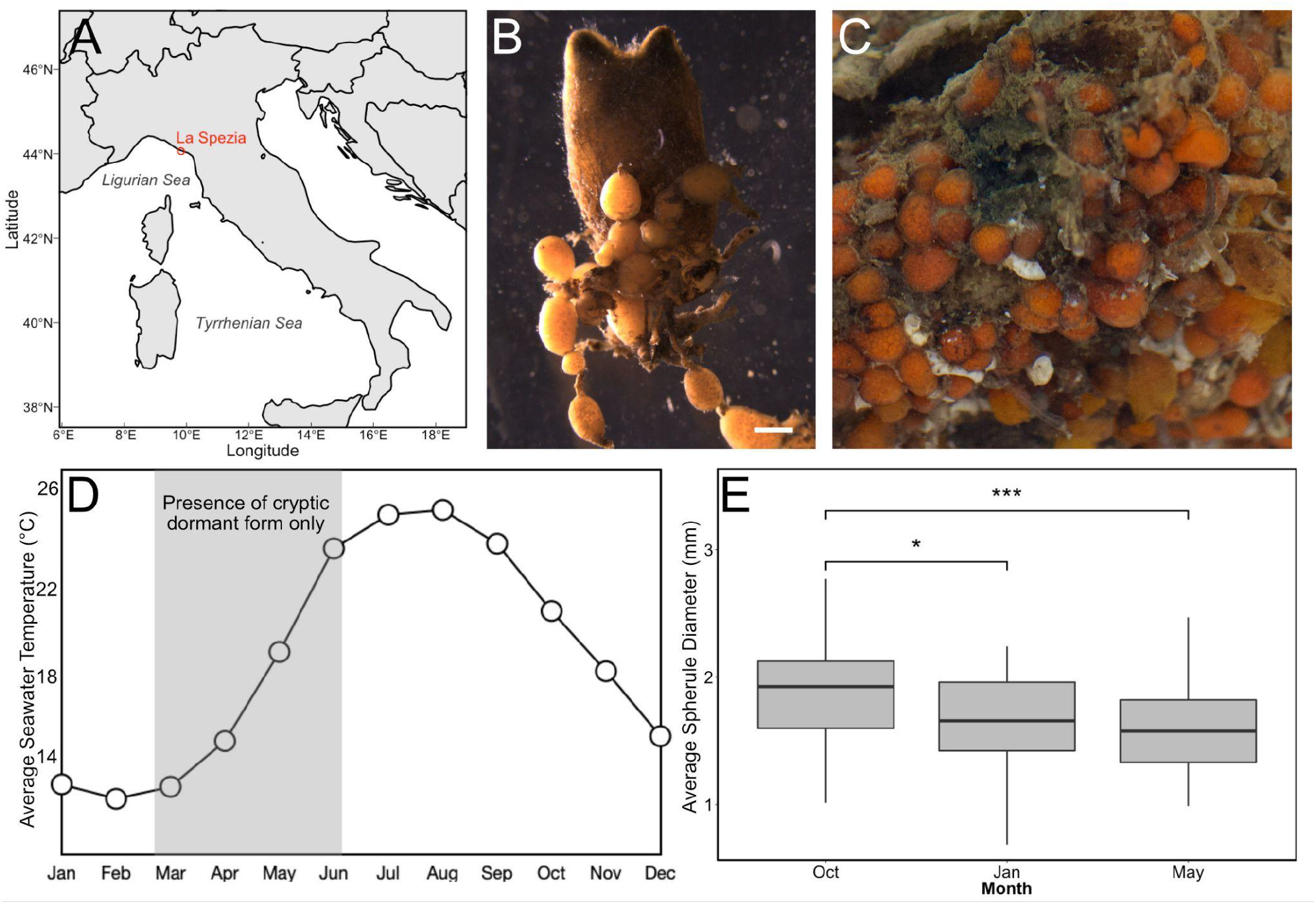
Field colony dynamics of *Polyandrocarpa zorritensis* in La Spezia, Italy. (A) Map shows collecting location in La Spezia in the NW of Italy. (B) Zooid collected from the harbor of La Spezia, with spherules attached along stolonal projections. Scale bar is 2mm. (C) Close-up of dormant spherules. (D) Average surface seawater temperature in La Spezia (NOAA Sea Surface Temperature satellite data from AVHRR Pathfinder SST). Shaded area indicates the period that animals exist in the cryptic dormant form. The rest of the year, both zooids and spherules are found on the ropes in the harbor. (E) Graph showing the change in average spherule size (n=45) over the year, with the largest spherules found in October, and the smallest spherules in May.

Combining our observations with the monthly temperature ranges of seawater in La Spezia from NOAA Sea Surface Temperature satellite data (AVHRR Pathfinder SST; Kilpatrick et al 2001) the largest spherules were obtained in the coldest months, right before the zooidal degeneration that occured in spring, when the water temperatures started to increase (Figure 5D). This suggests that during the cold season, the spherules accumulate nutrients and increase in size, preparing for survival after zooidal regression.

### 2. *Temperature change regulates the entry into and exit from dormancy in* P. zorritensis *and* C. lepadiformis

To test some possible abiotic environmental parameters influencing entry and exit of dormancy, small colonies of field-collected *P. zorritensis* (comprising zooids, budding nests, developing buds, and stolons) were maintained in controlled conditions in a flow-through aquarium system at 24°C and fed with a diet of living and concentrated algae. In order to determine if temperature would induce the production of dormant spherules, we removed a few small colonies (number of zooids=17; number of stolons=24; number of developing nests=5; number of non-abscissed nests=5) from the aquarium system and placed them at lower temperatures (10°C). Over the course of two months, we observed the production of dormant spherules. The zooids, when still alive, showed closed siphons and a low response to mechanical stimulation (n=15). The buds that we experimentally detached (abscised) from their “parent” zooid, which had already started the budding process, arrested development and degenerated (n=5). Stolonal growth continued, allowing the growth of the present budding nests (n=5) and the development of new ones (n=5). These structures were externally similar to spherules; they were dome-shaped and the tunic was so thick that it obscured internal vascular ampullae. Histological and ultrastructural observations were performed, showing that these structures were the same as spherules collected from the field (Scelzo et al. 2019). This represented the first successful production of dormant-stage specimens of this species in laboratory conditions.

To identify the colony tissue sensitivity and the precise temperature that triggered dormancy in *P. zorritensis*, we exposed the following groups-1) spherules collected in the field (n=10) stored at 10°C, 2) zooids with spherules (n=3) produced in laboratory after transfer at 10°C, and 3) budding nests (n=3) produced in the laboratory after germination of spherules and stored at 24°C- to four different temperatures (8°C, 12°C, 17°C, 22°C) and documented their development for 6 weeks. At the lowest temperature (8°C) all the zooids died while the spherules and the budding nests did not germinate. At 12°C, half of the zooids died or detached from the glass slide; stolonal growth was arrested; and new budding nests or developing buds did not form after 6 weeks. At this temperature all the spherules (collected in the field and produced in the laboratory) were activated and produced new zooids in 30-35 days. At 17°C the zooids stayed alive and well-attached to the glass slide, all spherules germinated (after about 20 days) and new stolons, budding nests, and spherules were produced. At 22°C the spherules and budding nests were rapidly activated (after two weeks new zooids were visible). However, half of the zooids died or were detached and few new stolons and nests were produced. These results suggested that the budding nests transferred to 8°C, 12°C, 17°C could form spherules. When the spherules (collected in the field or produced in the laboratory, both stored at 10°C) were transferred to lower temperatures (from 10°C to 8°C) they did not activate; however, when they were transferred to slightly higher temperatures (from 10°C to 12°C) germination was activated and new zooids were generated. Spherule germination was faster at higher temperatures (data not shown).

The ability to produce dormant forms under laboratory conditions was tested also in *C. lepadiformis*. Specimens of *C. lepadiformis* were collected at the same site as *P. zorritensis* and were cultured on glass slides at 18°C. As previously described by other authors (Giard and Caullery 1896, Della Valle 1915, Huxley 1926) the zooids emitted vascular stolons as prolongations of the postabdomen (Figure 4B). The stolonal tip sometimes started to bulge forming one or more lobes, which looked more pale and whitish than the rest of the stolon (Figure 4C). These structures represent the budding chambers, in which the budding process takes place once the hemocyte circulation is interrupted between the zooid and the budding chamber. At this temperature, other structures similar to budding chambers but more compact and whitish are produced along the stolon; they represent the winter buds, which are able to resist cold temperature according to other authors (Figure 4D; Berrill, 1951; Berrill and Cohen, 1936). Our results showed that the production by *C. lepadiformis* dormant forms, the winter buds, occurred at 18°C in laboratory conditions; and that similarly to budding chambers, the winter buds could be activated by isolation from the stolon. According to the above-mentioned authors, the milky coloration of winter buds is due to accumulation of trophocytes, i.e., hemocytes specialized for storage and transport of nutrients.

### 3. *Viability and range of resistance of* P. zorritensis *spherules and* C. lepadiformis *winter buds*

In order to determine the ability of dormant forms to resist sudden environmental changes, we exposed spherules and adult zooids of *P. zorritensis* to abrupt shifts in temperature (−20°C to 37°C) and salinity (10-44ppt) for 24 hours. Following the stress treatment, the specimens were transferred to 24°C and the viability of zooids and the germination of dormant forms were observed after two weeks. *P. zorritensis* zooids survived a minimal range of conditions (between 18 and 28°C and between 30-38 ppt), while spherules resisted a wider range of conditions (temperature fluctuation between 0 and 32°C and salinity between 15 and 45 ppt) (Figure 6).

**Figure 6:**
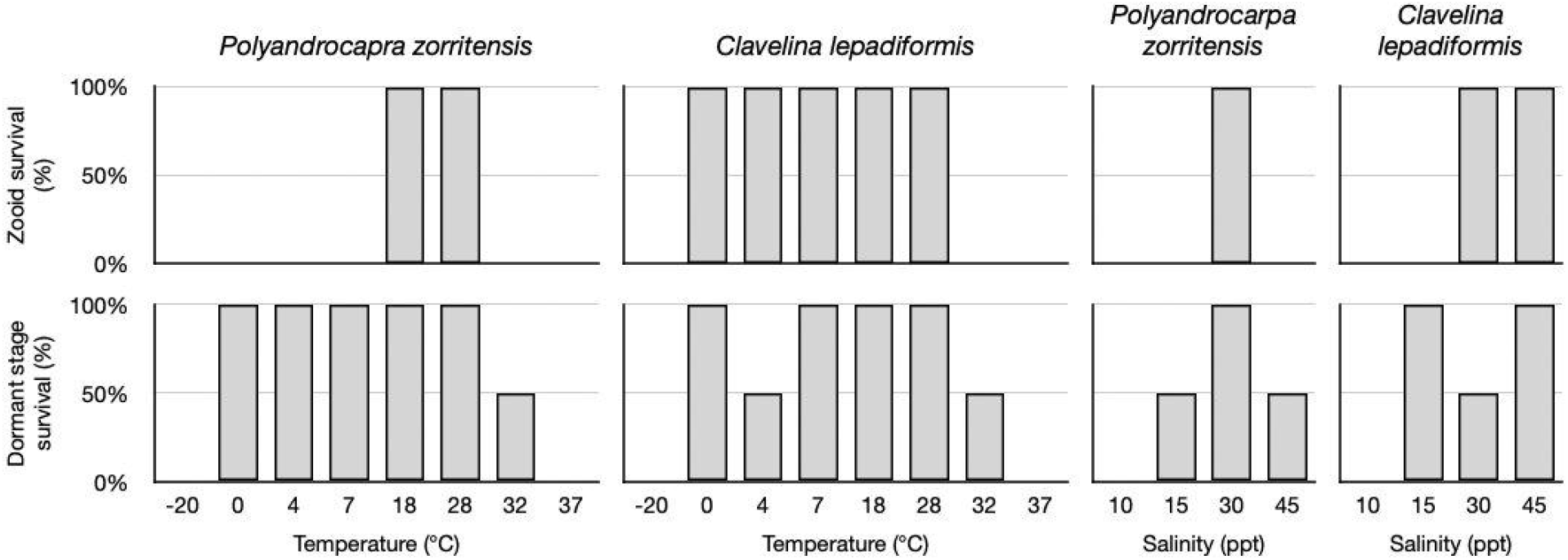
Dormant stages have broader capacity for survival, compared to zooids, in response to environmental stressors. Graphs of percent survival after 48h exposure to the indicated temperatures and salinity (n=2 per treatment) for *P. zorritensis* and *C. lepadiformis*.Top row shows survival rates of zooids. Bottom row shows survival rates for dormant structures (spherules for *Polyandrocarpa zooirtensis* and winter buds for *Clavelina lepadiformis*).

We repeated the same experiment with winter buds and *C. lepadiformis* zooids. Both stages were exposed for 24h to a wide range of temperatures (−20°C to 37°C) and water salinity (10-44ppt). Then, the specimens were transferred to 24°C and the viability of zooids and the germination of dormant forms were observed after one week. *C. lepadiformis* zooids were more resistant to lower temperatures (between 0 and 28°C) and higher salinity (30-45 ppt) than *P. zorritensis*. Winter buds resisted a wider range of conditions compared to zooids (temperatures between 0 and 32°C and salinity between 15 and 45ppt), similar to *P. zorritensis* spherules (Figure 6).

### 4. *Morphology and transcriptomic signature of* P. zorritensis *spherules and* C. lepadiformis *winter buds show structures and genes involved in the maintenance of dormancy*

From a histological perspective, a spherule appears as a cluster of monolayered chambers, the ampullae, interconnected by vessels, all embedded in a thick extracellular matrix called tunic (Figure 7E and 8A; Scelzo et al 2019). Tunic fibers of spherules are thicker than in the budding nests (∼20μm vs ∼5μm; Scelzo et al 2019). The epithelia of the ampullae is formed by cylindrical columnar cells of ∼10μm in length and ∼5μm in width (Figure 7F). The cytoplasm of the epithelial cells are filled with two types of electron-dense inclusions: small electron-dense granules of glycogen (0.5-1μm) and uniform electron-dense granules of lipids (2-5μm). The ampullae and vessels are filled with a heterogeneous population of hemocytes, among which it is possible to identify round undifferentiated cells of 5μm diameter (Figure 7H) which have a large nucleus-to-cytoplasm ratio, a prominent nucleolus, and with poorly developed endoplasmic reticulum and scarce mitochondria, reminiscent of ascidian circulatory stem cells referred to as hemoblasts (Kassmer et al 2020; Jiménez-Merino et al. 2019).

**Figure 7:**
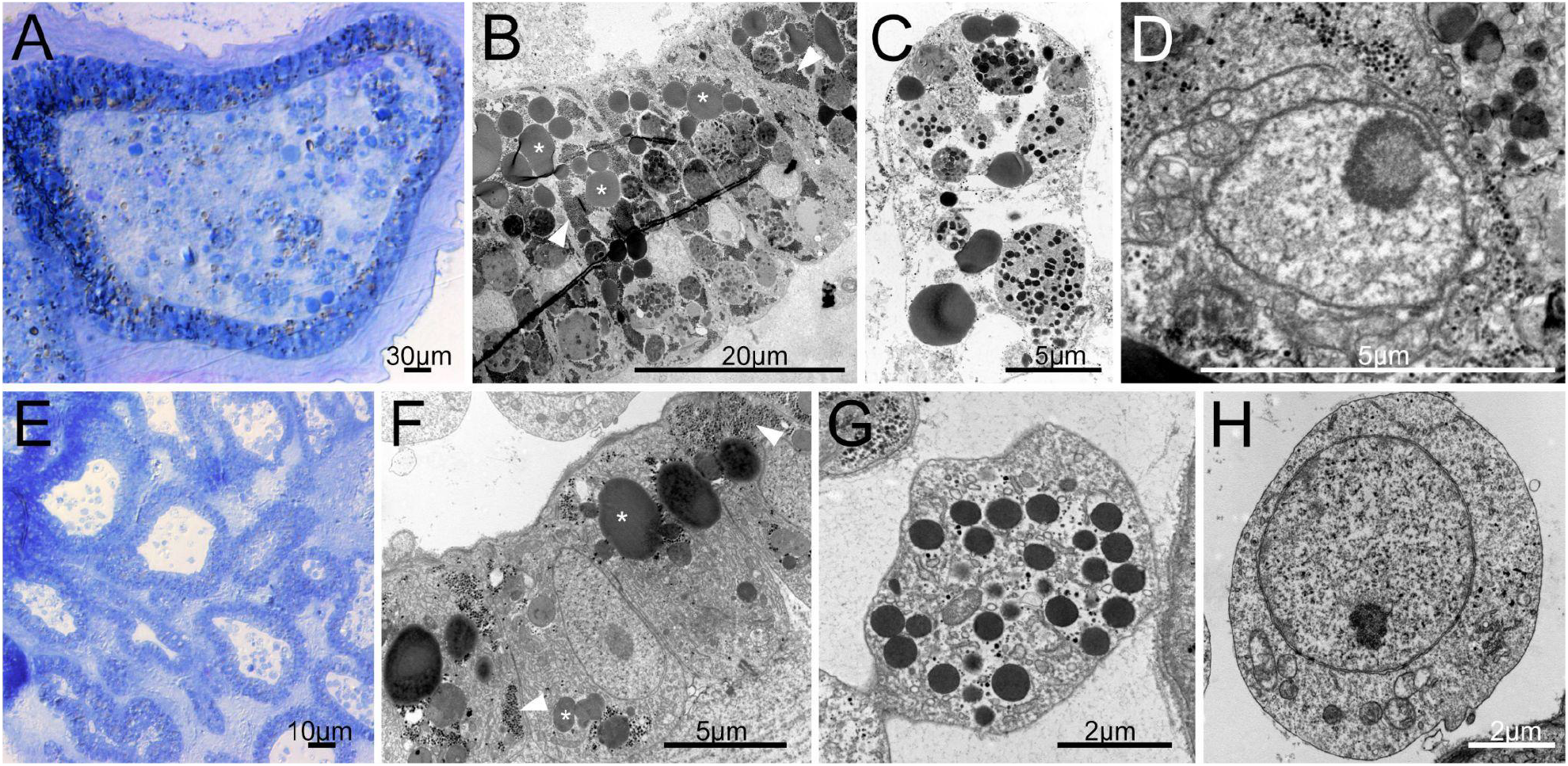
Cell types in dormant stages of *Clavelina lepadiformis* and *Polyandrocarpa zorritensis* include nutrient storage cells and putative stem cells. (A-D) *C. lepadiformis*: (A) Methylene blue staining of winter bud histological section shows a thick layer of epithelial cells surrounded by tunic and enclosing abundant mesenchymal cells. (B) TEM image of epithelial cells, many with lipid and glycogen inclusions, labeled with asterisk and white arrowheads, respectively. (C) TEM image of a phagocyte with contents inside mesenchymal space. (D) Putative hemoblast cell inside mesenchymal space, as indicated by large nucleolus and small cell size (about 5 microns). (E-H) *P. zorritensis*. (E) Methylene blue staining of spherule histological section shows numerous ampullary spaces lined with epithelium and filled with sparse hemocytes. (F) TEM close-up image of epithelial cells of ampulla, showing inclusions of lipids (asterisks) and glycogen (white arrowheads). (G) Example of a mesenchymal cell with inclusions. (H) TEM image of putative hemoblast cell within the mesenchymal space, as indicated by large nucleolus, small cell size, and round cell shape.

**Figure 8:**
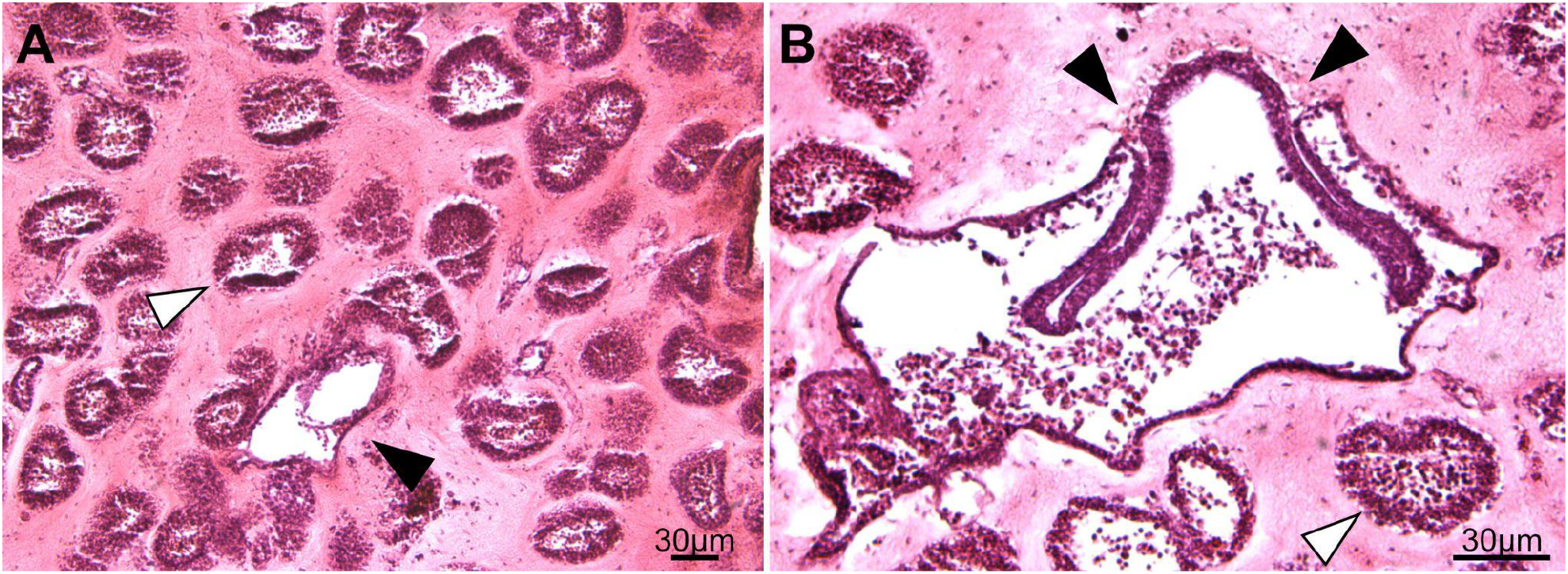
Germination transition in *P. zorritensis*. Transversal histological sections stained with hematoxylin and eosin of germinating spherule after 48 hours at 24°C. (A) The central vessel (black arrowhead) is distinguishable from the ampullary ramifications (one example marked with white arrowhead). (B) In a section through the center of the spherule, the vascular epidermis is invaginating (black arrowheads indicate location of the invaginations). White arrowhead indicates ampullary ramification.

In a *C. lepadiformis* winter bud, the epidermal cells around the vascular lumen are filled with electron-dense lipid inclusions in the cytoplasm—similarly to *P. zorritensis* spherules, which include small glycogen granules, homogeneous electron dense granules, and heterogeneous electron dense inclusions. The vascular lumen of the winter bud contains many more mesenchymal cells and more densely-packed cells than the stolonal lumen. The winter bud lumen contains large cells of more than 20μm that appear to be macrophages (Figure 7C). The phagosomes of the macrophage-like cells are filled with material containing electron-dense inclusions (Figure 7C). These cells are tightly packed and surrounded by non-cellular debris (Figure 7A-D).

In order to elucidate the molecular signature of dormancy, we examined gene expression across life stages in both *P. zorritensis* and *C. lepadiformis* (Figure 9). Of the orthologous DEGs that were up-regulated in dormant tissues in both species, 530 genes were shared. Among the shared up-regulated genes in dormancy, we examined KEGG pathway enrichment and found an over-representation of genes in a number of pathways, such as the HIF-α pathway and the insulin signaling pathway (Figure 10). Key members of both of these pathways are upregulated in both species (KEGG diagrams in Supplementary Figure 1).

**Figure 9:**
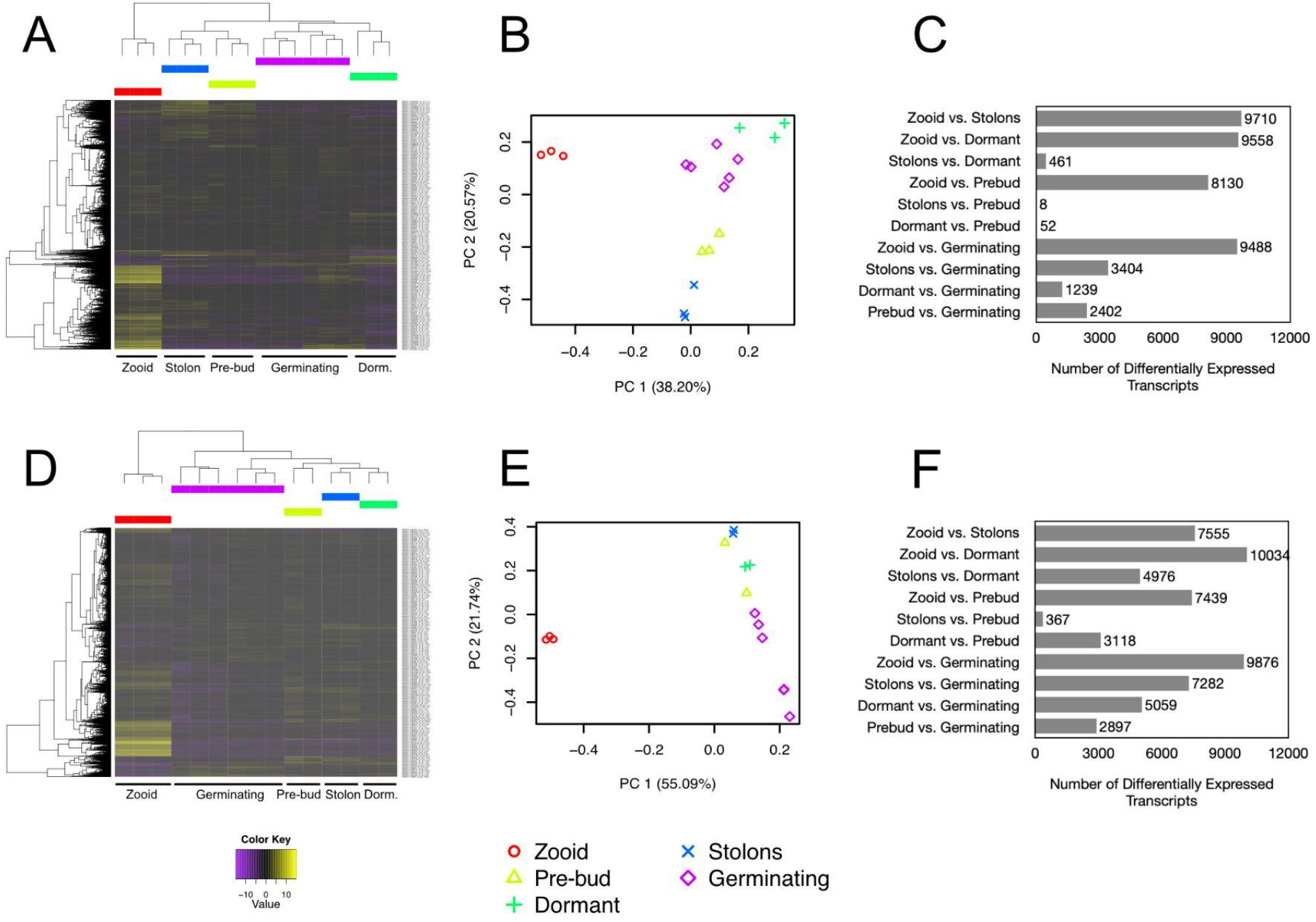
Differentially expressed transcripts during life stages in *Polyandrocarpa zorritensis* and *Clavelina lepadiformis*. Heatmaps of differentially expressed transcripts across samples: (A) *P. zorritensis* and (D) *C. lepadiformis*; each row represents expression value of a single transcript as shown in the color key below plots, violet indicates low expression and yellow indicates high expression; columns show clustered life stage replicates, bars in red=zooid, blue=stolon, yellow=pre-bud, green=dormant, and magenta=germinating stage. Principal Component Analysis showing relationships among and across samples: (B) *P. zorritensis* and (E) *C. lepadiformis*. Key below indicates colored symbols for each life stage sample type. Number of differentially expressed transcripts in each pairwise comparison between sample types for each species: (C) *P. zorritensis* and (F) *C. lepadiformis*.

**Figure 10:**
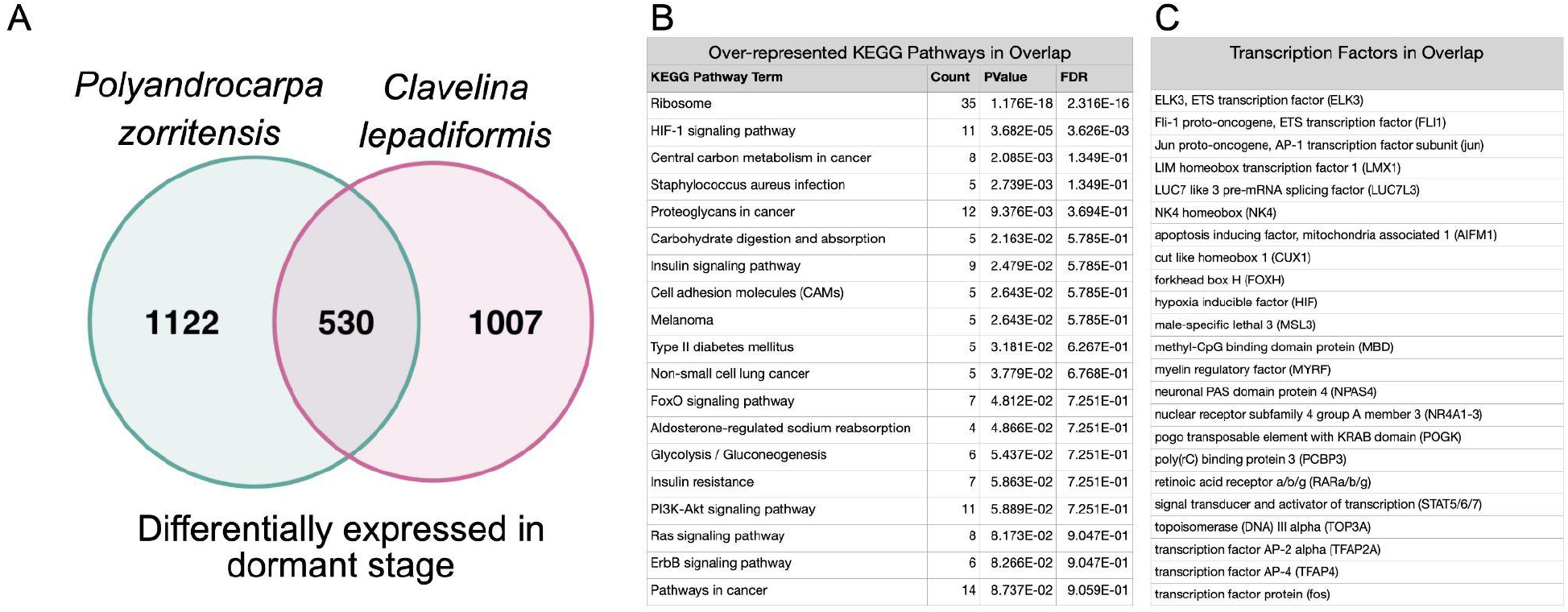
Shared up-regulated transcripts and pathways during dormancy in *P. zorritensis* and *C. lepadiformis*. (A) Venn diagram showing the number of orthologous upregulated transcripts during dormancy in *P. zorritensis* and *C. lepadiformis*. (B) KEGG pathway terms enriched in the overlapping set of 530 transcripts upregulated in both species. (C) Transcription factors upregulated in dormancy in both species

We also examined upregulated transcription factors in dormancy that were either shared between species or found exclusively in one species (Figure 10C). Some of the shared upregulated transcription factors were involved in HIF-α signaling, such as a hypoxia inducible factor alpha (HIF-α), and in insulin signaling. Additionally, we found shared up-regulation of a number of genes implicated in the modulation of stem cell maintenance, proliferation, and differentiation. These included: orthologues of AP-1 transcription factor (fos and jun), NK4 homeobox, nuclear receptor subfamily 4 group A (NR4A1-3), retinoic acid receptor beta (RAR-A/B/G), and signal transducer and activator of transcription (STAT5-7). Some of the shared transcription factors upregulated in dormancy have gene silencing or chromatin remodeling functions, such as methyl-CpG binding domain protein 2 (MBD2-4) and male-specific lethal 3 (MSL3) (Du et al 2015; Rea and Akhtar 2006).

### 5. Exit from dormancy shares morphological/molecular features as in whole body regeneration or budding in colonial ascidians

In order to test if the developmental events in spherule germination are comparable to budding in *P. zorritensis* budding nests, spherules maintained at 10°C were transferred to a higher temperature (24°C) and fixed after 48 hours. In the developing spherules, the vascular epidermis started to invaginate in the same way as occurs in developing budding nests (Figure 8B). These results suggest that the morphogenetic epidermal movements occurring in germinating spherules are comparable to the vasal budding processes described in the development of budding nests (Scelzo et al 2019).

We found that 674 genes are up-regulated during germination in both species (Figure 11A). KEGG pathways upregulated in this shared set of genes include many metabolic pathways (Figure 11B). We also found a number of shared up-regulated transcription factors. This includes developmental genes such as NK4, grainyhead-like transcription factor (GRHL1/2/3), and a distal-less homeobox (DLX1/6), and retinoic acid receptor beta (RAR-A/B/G) (Figure 11C). Interestingly, some transcription factors that showed upregulation in germination are also genes that we found to be upregulated in dormancy, including NK4, RAR-A/B/G, and forkhead box H (FOXH).

**Figure 11:**
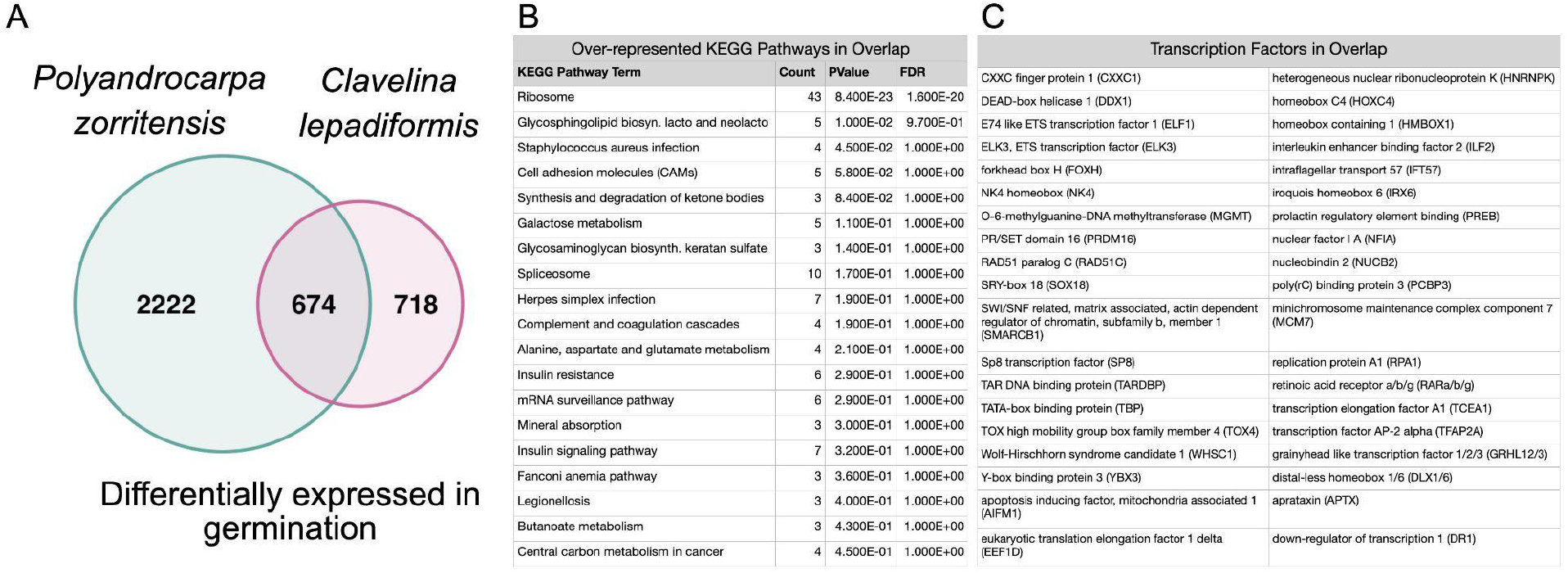
Shared up-regulated transcripts and pathways during germination in *P. zorritensis* and *C. lepadiformis*. (A) Venn diagram showing the number of orthologous upregulated transcripts during germination in *P. zorritensis* and *C. lepadiformis*. (B) KEGG pathway terms enriched in the overlapping set of 674 transcripts upregulated in both species. (C) Transcription factors upregulated in germination in both species.

## Discussion

### Ecological consequences for dormancy in clonal species

We report the disappearance of zooids of *Polyandrocarpa zorritensis* at the end of the winter in NW Italy, with the maintenance of only a cryptic dormant form: small spherules hidden under epibionts. Such small, relatively undifferentiated, and practically unrecognizable structures remain dormant for months and have the ability to reconstitute a thriving population of differentiated feeding animals when conditions improve. By the end of the spring, through a mass-scale regeneration event, spherules are able to restore the entire population of filter-feeding colonies. Our data on spherule diameter over the year suggests that spherules get smaller as the zooid die-off period approaches, which may be due to the nutritive and energetic demands of cells within the spherules over the year. Winter buds of *Clavelina lepadiformis* have also been found to exist in the absence of zooids. Winter buds without zooids have been documented in the North Sea in the winter months (Orton 1914, 1921), in the summer months in the eastern Mediterranean, and both *C. lepadiformis* and *C. gemmae* regress during summer months (called aestivation) in the western Mediterranean (Mukai 1977; de Caralt 2002; Turon 2005). These results show that the dormant stages of both *C. lepadiformis* and *P. zorritensis* are resistant to extremes in temperature and salinity. Thus, this linked capacity for dormancy and regeneration is likely an important adaptation that allows these species to survive drastic seasonal changes.

Our field observations suggest that even in conditions favorable for budding, the dormant forms are continuously produced as stolon mats under the colony and are ready to assure survival if conditions become adverse. In both species, the pre-buds and dormant structures are produced throughout the time that the filter-feeding zooids are also present. As soon as these structures detach from the parent zooid, budding is triggered. However, if the stolonal connection is maintained, the pre-buds continue to grow in size, acquiring additional nutrients, and budding is inhibited. This occurs once the zooids have propagated to take up significant space such as to limit available surface area for further zooid production. When the zooids are highly packed together, the pre-buds are protected from abscission, and when they maintain the connection with the parent, they continue to grow into larger dormant structures. The connection with the parent is also maintained more readily at low temperatures in laboratory conditions.

The capacity for dormancy followed by regenerative germination allows for survival over time periods of unfavorable conditions, but this adaptation likely allows for survival across space and into new habitats. Both *P. zorritensis* and *C. lepadiformis* are well-documented invasive species in numerous regions of the world. *C. lepadiformis*, which is a native of the Mediterranean Sea, Atlantic subarctic regions (i.e., Norwegian Sea and Greenland), and Bay of Biscay, has successfully spread to the Atlantic coasts of North and Central America, and Brazil (Pyo and Shin 2011; Reinhardt et al 2010; Turon et al 2003). *P. zorritensis*, described for the first time in Peru (Van Name, 1931), has been reported in the last decades at sites all over the world, especially in harbors (Carman et al 2011; Lambert and Lambert 2003). In the Mediterranean Sea, *P. zorritensis* has been recorded in Italy (Brunetti, 1978; Brunetti and Mastrototaro 2004; Mastrototaro et al 2008; Stabili et al 2015), Spain (Turon and Becerro 1992) and very recently in France, in the Thau pond (reported on the DORIS website https://doris.ffessm.fr/Especes/Polyandrocarpa-zorritensis-Polyandrocarpe-de-Zorritos-5004).

Tolerance to a range of environmental conditions has been proposed to help account for its invasive success. The physiological responses to abiotic parameters such as light, pressure, and salinity have been investigated in the larvae of *P. zorritensis* (Sumida et al 2015; Vázquez and Young, 1996, 1998) but these results were insufficient to explain the rapid expansion of this species. Our data show that *P. zorritensis* spherules are capable of resisting a wide range of environmental extremes, and these small structures can be easily transported and reestablish a new colony. Thus, *P. zorritensis* invasions in semi-enclosed basins with peculiar abiotic conditions, like the case of Taranto Sea in southern Italy (Mastrototaro et al. 2008), may have been facilitated by the resistance of the spherules to large changes in salinity and temperature. Overall, the unique dormant phase of the life cycle and the ability to survive in a non-feeding state may underline the extreme success in invasion ability of these species.

### Convergences in cellular makeup of dormant structures

The dormant structures in both *C. lepadiformis* and *P. zorritensis* contain a relatively small amount of cell morphotypes. Those include cells that store the nutrients for the maintenance of the dormant stage, cells that have the potential to contribute to the development of new buds (undifferentiated mesenchymal cells), and cells with protective function, including tunic-secreting cells. In many dormant tunicates, specialized mesenchymal cells called trophocytes, storage cells, or granular amebocytes, accumulate nutritive reserves that are used during the development of the new zooid (Brien and Brien-Gavage 1927; Mukai et al 1983; Fujimoto and Wantatabe 1976; Cima et al 2016; Hyams et al 2017). In *C. lepadiformis*, the trophocytes accumulate reserves by sequestering breakdown products of dying and degenerating zooids (Kreb 1908; Driesh 1902; Shultz 1907). While a few authors speculated that these cells represent the undifferentiated regenerative cells (Orton 1921; Spek 1927), most agree that they correspond to large phagocytic migratory cells that contain cytoplasmic glycoproteins and other inclusions, but have no direct role during the differentiation occurring during germination (Brien 1930; Reis 1937; Fisher 1937). Fisher (1937) also documented “cell excretions”, which may be decaying cell material that is excreted into the vasculature to be used for nutrition of the developing bud. Fujimoto and Wanatabe (1976) examined budding tissues in *Polyzoa vesiculiphora* and found the occurrence of granular amebocytes that contained glycogen particles, lipid droplets, large protein granules, and autophagosomes, which they classified as trophocytes. They documented that these cells were phagocytized over the course of bud development, then disappeared, suggesting that they are important for nutrient supply to the bud. In our ultrastructural analysis of *C. lepadiformis* winter buds, we documented large cells filled with phagosomes and inclusions of glycogen and lipids, which we also believe represent the trophocytes. We also found lipid and glycogen-rich cells present in the epithelia of the winter bud in *C. lepadiformis*. In contrast, we did not find any mesenchymal trophocytes in *P. zorritensis* that could be used for nutrition, and therefore we propose here an alternative strategy this species may use to sequester nutrients within the dormant form. The vascular epithelium of *P. zorritensis*, unlike *C. lepadiformis*, is highly ramified within the spherule, providing many epithelial cells, filled with nutrient inclusions of lipids and glycogen, that may be used during dormancy and to support germination. Thus, the two species have acquired unique adaptations for endowing the dormant form with distinct supplies of nutrition to last through the dormant period and during germination.

The pluripotent cells or tissues that give rise to the mesodermal layer (the mesentoblast or “inner vesicle” of the characteristic double-vesicle stage of bud development) differs across diverse tunicate species. Early work showed that the inner vesicle in *C. lepadiformis* arises from cells from the stolonal septum (Seelinger 1882; Van Beneden and Julin 1886; Giard and Caullery 1896; Garstang 1928; Brien and Brien-Gavage 1928; Brien 1930), whereas Fisher (1937) cultured winter bud cells and found that the small undifferentiated cells contributed to the formation of new budding tissue. Our ultrastructural examination of winter buds lead to the identification of small cells with a prominent nucleolus, which likely represent the undifferentiated cells that give rise to the new buds. However, we could not clearly visualize the stolonal septum within the winter bud. So, how the septum and undifferentiated cells play a part in the budding process in both normal buds and winter buds is still unclear. *P. zorritensis*, on the other hand, lacks a stolonal septum, and the source of the inner vesicle in budding nests is known to be the vascular epithelium (Scelzo et al 2019). However, we did observe mesenchymal putative stem cells (hemoblasts) in the vasculature of *P. zorritensis* during dormancy. A role for hemoblast cells during germination is possible, but has not been documented.

### Shared molecular mechanisms of dormancy across distantly related tunicates

Our analysis involves two phylogenetically distant tunicate species that most likely have independently acquired dormancy. Both species express a number of common genes and pathways during the dormant phase. For example, both species show up-regulation of key components of the HIF1-α signaling pathway and the insulin signaling pathway. HIF-α signaling is utilized in other species during dormancy (Dias et al, 2021). HIF-α signaling induces a metabolic rewiring that promotes anaerobic glycolysis and a shift from glucose to fatty acid combustion that helps maintain cellular quiescence (Dias et al., 2021). Further, the insulin signaling pathway has been shown to be key to the control of diapause in some insects (Han and Denlinger 2011), as well as the dauer diapause stage in nematodes (Kimura et al. 1997; Ogawa and Brown, 2015). In low nutrient situations, insulin signaling is reduced, which activates FoxO/DAF-16, which lowers the production of ROS to protect against oxidative damage, and regulates both the formation and morphogenesis of the diapause stage (Dias et al 2021; Ogawa and Brown 2015). These mechanisms have also been described during mammalian hibernation and cellular quiescence in hematopoietic stem cells (Dias et al 2021). Our findings further support the occurrence of a core program of dormancy in animals that is related to the physiological and developmental demands of this unique stage.

Among the upregulated genes during dormancy in both *P. zorritensis* and *C. lepadiformis* we found transcription factor AP-1 (activating protein-1), which is known to regulate gene expression in response to various stimuli, such as stress, hypoxia, or growth factors (Wisdom 1999). The AP-1 family of transcription factors contains homodimers and heterodimers of fos and jun subunits, among others (Karin et al 1997). We find fos and jun up-regulation in dormancy in the two tunicate species we examined. AP-1 is known to have a role in regulation of cellular proliferation, senescence, and differentiation, including in the context of regeneration, where these genes are often some of the first responders after injury (Karin et al 1997; Srivastava, 2021). Thus, AP-1 may play an important role in maintaining cells in a non-proliferative state or undifferentiated state during dormancy. Sensing changes in environmental conditions is key for sessile organisms to make decisions about resources committed to growth and development. As oxygen levels and nutrient availability are two of the most variable environmental parameters, it is not surprising that both HIF, insulin signaling pathways, as well as quick-responding genes such as AP-1 family factors would be shared regulators of dormancy. In the future, higher resolution lab studies with precisely manipulated food and oxygen levels could be used to compare responses between species, and identify the order of molecular events.

### Regenerative emergence (germination)

In both *P. zorritensis* and *C. lepadiformis*, the dormant state appears to be overlayed onto the pre-budding stage of the life cycle. As dormancy ends, and germination begins, the budding process resumes. We found a number of transcription factors that are highly expressed in this germination stage that are shared between both species. For instance, NK4 (the tunicate homologue of vertebrate Nkx 2.3/5/6) was previously found expressed at the onset of two independently evolved types of budding, vasal budding in *Polyandrocarpa zorritensis* and peribranchial budding in *Botryllus schlosseri* (Alié et al. 2018). NK4 is also expressed in various embryonic and larval territories and notably required for heart formation (Wang et al. 2013, Prünster et al. 2019). The high expression of NK4 during dormancy exit may indicate another pleiotropic role of this transcription factor, but it may also reflect the homology between dormant and asexually reproducing structures.

We found that a retinoic acid receptor is upregulated in both dormancy and during germination in both species. Additionally, the enzymes retinal dehydrogenase and retinol dehydrogenase show upregulation during germination of *P. zorritensis* and *C. lepadiformis* respectively (Supplementary Figure 2). Together these results are suggestive of a role for retinoic acid (RA) signaling during germination. RA signaling pathway is implicated in budding in *Polyandrocarpa misakiensis* (Kawamura et al 2013) and whole body regeneration in *Botrylloides leachii* (Rinkevich et al 2007a). In our study, we only examined the expression at a single time point during germination, and it is possible that sequence data from more time points during this process would allow for better resolution of the gene expression dynamics. However, even with this single snapshot, finding shared upregulated genes in germination across distantly related clonal tunicates suggests that these species may employ similar mechanisms in their asexual budding processes. As these species have evolved budding independently from each other, the use of RA signaling in bud formation, if it indeed has a role, would likely represent convergent co-option from its role in embryonic development.

### Conclusions

While the origin of a novel dormant life stage appears to have evolved independently multiple times among asexually replicating tunciates, we have documented a number of ecological, morphological, and cellular convergences across two separate origins of dormancy. In both species, the animal produces a modified stolon filled with nutritive cells and potentially multi/pluripotent cells, which can withstand stressful environments that the adult zooids cannot survive, and is linked to sensing mechanisms to adjust metabolism and time the initiation of budding to when conditions improve. In these cases, dormancy utilizes some of the same cellular and morphological aspects of the budding process that itself evolved independently in each clade, such as the use of the stolon tissues and cells. Our results point to the ways that species reuse existing genes and pathways for the origins of budding and dormancy in tunicates. In particular, our results suggest: (1) conserved eco-physiological sensing mechanisms (e.g., HIF-α signaling) are repeatedly co-opted for the origins of dormancy, as has been found in numerous animals (Wilsterman et al 2021), and (2) conserved developmental/regenerative mechanisms (e.g., RA signaling and NK4) are repeatedly co-opted for the origins of budding/germination.

## Material and methods

### Sample collection and animal husbandry

Colonies of *Polyandrocarpa zorritensis* and *Clavelina lepadiformis* were collected in the harbor of La Spezia, Italy (Assonautica Benedetti, 44°06′10.7′′N 9°49′34.5′′E). Colonies and spherules of *P. zorritensis* were maintained in the laboratory as reported in Scelzo et al. (2019). In order to determine spherule sizes over the course of a year, *P. zorritensis* were collected on three different docks of the marina “Porto Mirabello” during fall (October 2018), winter (January 2019), and spring (May 2019) and spherule diameters was measured. Spherules were collected from the same rope (at marina slips C2, G7, and E15) at each time point with the goal of following the size of spherules of a single colony over time (n=15 per rope per timepoint; n=45 per timepoint). Diameters (along the longest axis) were calculated from microphotographs in ImageJ. Average spherule diameters were compared using a repeated measures ANOVA conducted in R. Colonies of *C. lepadiformis* were maintained at 18°C in a closed sea water system and fed with a mixture of living algae (*Tisochrysis lutea* and *Chaetoceros*) and concentrated algae (Isochrysis 100 and Shellfish Diet 1800, Reed Mariculture Inc, Campbell, CA, USA). *C. lepadiformis* zooids were attached to glass microscope slides with sewing thread and allowed to adhere by stolonal growth at 18°C for one week. The thread was then removed and the animal was allowed to continue growing until use in experiments. Slides were cleaned twice weekly under a dissecting microscope to remove biofilm and epibiont growth. Microphotographs of animal morphological stages were taken on a Leica M165FC stereoscope.

### Light and transmission electron microscopy (TEM)

Inclusion of *P. zorritensis* specimens for paraffin sectioning and hematoxylin/eosin staining were performed as described in Alié et al. (2018). Samples of *P. zorritensis* and *C. lepadiformis* for semi-thin and ultra-thin sectioning were fixed, embedded and treated as reported in Scelzo et al. (2019).

### Effects of environment on dormancy of P. zorritensis and C. lepadiformis

Adult zooids and dormant stage animals were maintained in incubators at four constant temperatures: 8°C, 12°C, 17°C, 24°C. In each condition, animals were maintained in aquarium tanks filled with ∼700mL sea water with constant aeration and fed three times a week with a mix of living algae (*Tisochrysis lutea* and *Chaetoceros*) and concentrated algae (Isochrysis 100 and Shellfish Diet 1800, Red Mariculture Inc). Water changes and animal cleaning were carried out two times a week.

To test the temperature resistance range of zooids and dormancy stages in both species, the specimens were transferred to 50mL Falcon tubes (Corning, Glendale, Arizona) in filtered sea water and exposed to different temperatures in incubators (4°C, 18°C, 28°C, 32°C, 37°C) or on ice (for the 0°C condition) for 24 hours. Then, the *P. zorritensis* and *C. lepadiformis* specimens were gradually transferred respectively to 24°C and 18°C. For the following two weeks, their survival or budding was observed and they were cleaned and fed as in the non-treated specimens.

To test the salinity resistance range of zooids and dormancy stages in both species, animals were exposed to 10 ppt, 15 ppt, 30 ppt, 39 ppt, and 45 ppt salinity water (n=2 for each treatment and each stage). The low salinity water conditions (10, 15, and 30 ppt) were prepared by adding deionized water to seawater. The high salinity water was prepared by adding artificial sea water to filtered seawater (Red Sea Salt, Red Sea). For each condition, the specimens were transferred to a 50mL Falcon tube (Corning, Glendale, Arizona) and incubated at 24°C (*P. zorritensis*) or 18°C (*C. lepadiformis*) for 24 hours. Then, they were gradually transferred to filtered sea water at 39 ppt and observed for the following two weeks.

### Differential gene expression analysis

Five life stages were chosen for RNA sequencing of each species: zooid, stolon, prebud (budding nest/chamber), dormant stage, and germination. For each stage, three replicate samples were prepared for sequencing. For each stage, RNA was extracted from three replicate samples. Tissue from three or more individuals was pooled to obtain enough RNA for sequencing. For the dormant stage *P. zorritensis*, spherules were collected from the field in the spring (the time of the year in which only dormant stages were present) and the tissue was immediately flash-frozen. Additional material from the same rope was transported back to the lab, and germinated to generate new zooids, stolons, and buds, from which we extracted RNA for those stages. For *C. lepadiformis*, we extracted all the material from the same colony in the lab. The “germination” samples were germinated for 48 hours, with 3 replicates at 18°C and 24°C each. The replicates were combined during analysis.

RNA was extracted with the NucleoSpin RNA extraction kit (Machery-Nagel) and sample quality and quantity were verified with the B2100 BioAnalyzer Instrument and Bioanalyzer RNA 6000 Nano assay (Agilent Technologies, Inc., Santa Clara, CA, USA). Frozen RNA was sent for cDNA library preparation (using NEB Next® Ultra RNA Library Prep Kit for Illumina) and Illumina paired-end 150bp sequencing at Novogene Bioinformatics Technology Co. Ltd (Tianjing, China). Sequencing was done on a Novaseq 6000 PE150 Illumina sequencing platform. Samples were split between two sequencing lanes with at least one of each replicate per stage in each lane to reduce lane-effects.

Raw sequencing reads (541,854,284 for *P. zorritensis* and 521,595,519 for *C. lepadiformis*) were cleaned and trimmed. First, the program Rcorrector was used to correct random sequencing errors and to eliminate read pairs where reads were flagged as ‘uncorrectable’ (Song and Florea, 2015). Next, Trim Galore! (Krueger, 2015) was used to remove low-quality reads from the dataset (with a quality Phred score cutoff of 5, a minimum read length threshold of 36 bp, a stringency parameter of 1 for adapter sequence overlap, and a maximum allowed error rate of 0.1) and to trim Illumina adapters sequences from raw sequencing reads (https://github.com/FelixKrueger/TrimGalore; Martin, 2011). Ribosomal RNA was removed by eliminating reads that mapped to the SILVA large and small subunit databases (SILVA_132_LSUParc_tax_silva.fasta and SILVA_138_SSURef_NR99_tax_silva.fasta) using Bowtie2 (Langmead and Salzberg, 2012). Finally, we used the software FastQC to identify overrepresented sequences from the read set, which we removed (Andrews, 2010). After the preprocessing steps, 496,811,038 reads remained for *P. zorritensis* and 506,081,729 reads remained for *C. lepadiformis*.

Transcriptomes were assembled using the software Trinity with *in silico* read normalization (Grabherr et al., 2011). For each species, read data after the above preprocessing steps were pooled across tissue types to create a single reference assembly per species. The reference assemblies (809,462 and 492,887 contigs for *P. zorritensis* and *C. lepadiformis*, respectively) were taxon-filtered to remove contaminant transcripts. First, two BLAST searches were conducted to identify possible contaminants: a Diamond Blastx against the uniprot ref database and a blastn search against the NCBI’s nr database (both with E-value set to 1e-25). The program Blobtools (Laetsch and Blaxter 2017) was used to assign a taxon to each transcript using the blast results (using option “bestsumorder”) and then to remove transcripts that were assigned to anything other than chordate, no-hit, or undefined. Proportion of contaminants was 12.5% for *P. zorritensis* and 18.3% for *C. lepadiformis* Next, the program Transdecoder (https://github.com/TransDecoder/TransDecoder) was used to select only the single best open reading frame per transcript, and only keep those with a minimum length of 90. The program cd-hit (Li and Godzik 2006) was used to collapse similar transcripts (parameters: -M 0 -T 20). Low-expression transcripts were removed using Kallisto to estimate abundance and the trinity script “filter_low_expr_transcripts.pl” to remove transcripts with expression level lower than 5 TPM. Final assemblies were 27,938 transcripts for *P. zorritensis* and 18,610 transcripts for *C. lepadiformis*. Assemblies were assessed by looking at basic summary matrices and by quantifying completeness using the program BUSCO (v3) by searching for a curated set of single copy orthologs present in all metazoans (metazoa odb9 database) (Seppey et al., 2019). BUSCO scores of final reference assemblies were: 68.0% for *P. zorritensis* and 70.7% for *C. lepadiformis*, suggesting that assemblies were relatively complete.

Genes were annotated first by DIAMOND blastx (Buchfink et al, 2021) to human refseq database (e-value 1e-25), then those without human ortholog matches, further DIAMOND blastx searches were conducted against the UniProt reference proteome database (e-value 1e-25) to identify lineage-specific transcripts (The UniProt Consortium, 2020). For *P. zorritensis*, 7,611 transcripts were human-annotated and 1,518 were annotated with non-human matches. For *C. lepadiformis*, 8,037 transcripts were human-annotated and 2,387 transcripts were annotated with non-human matches.

The program Kallisto (v. 0.46.0) was used to quantify abundances of transcripts and align preprocessed reads onto the assembled contigs (Bray 2016). This was done using scripts bundled with the Trinity package (Grabherr et al., 2011). Mapping rates were 54.2-60.8% for *P. zorritensis* and 60.5-70.9% for *C. lepadiformis*. The Trinity analysis program ‘PtR’ was used to compare biological replicates across samples, by constructing a correlation matrix for each sample, and principal components analysis (PCA) plots, labeled by stage and replicate. These analyses showed a good correlation between the replicates, and separation between tissue types (Figure 9).

Differential transcript expression was assessed using the program voom (Law et al., 2014) implemented within the Trinity package. All pairwise comparisons of sample type within species were tested. Transcripts were considered differentially expressed if they showed any difference in expression between sample types (C=0) with a false discovery rate (FDR) p-value of <0.001. Heatmaps of the differentially expressed transcripts were generated to cluster the transcripts according to their patterns of differential expression across the samples. Subclusters with genes of similar expression patterns were then calculated by cutting the heatmap by 30% percent tree height for *C. lepadiformis* and 50% tree height for *P. zorritensis*.

In order to identify genes that are consistently upregulated in each tissue/sample-type, we computed tissue-specific expression using perl scripts packaged with Trinity. We selected the “DE_pairwise_summary.txt.class_down_priority” files, which maximizes genes placed into the up-regulated category. Orthologous genes between the two species were identified using a reciprocal best blast hit strategy and the program Blast-Besties (https://github.com/Adamtaranto/blast-besties) using a e-value cutoff of 0.001, a length cutoff of 40 basepairs, and a bitscore cutoff of 100. This generated 6,137 orthologs. Of these orthologs, 4,345 were annotated with human reference data, and 826 did not have human matches but did have close matches in the uniprot proteome database, including many tunicate genes. From the orthologous subset between the two species, we determined the overlapping transcripts upregulated during dormancy and during germination in both species. The data was organized in R and Venn diagrams were produced using the package VennDiagram (v 1.6.0) (Chen and Boutros 2011). The shared annotated gene lists were used to determine the top over-represented KEGG pathways (Kanehisa and Goto 2000; Kenehisa, 2019; Kanehisa et al 2021). All annotated transcripts were used as background for the over-representation analysis calculations, which were done with the program DAVID (Huang et al 2009a and 2009b).

### Phylogenetic analysis

For thoses genes whose orthology has been confirmed, alignments were constructed by performing blastx using *P. zorritensis* or *C. lepadiformis* contigs as queries against *Homo sapiens, Mus musculus, Danio rerio*, Echinodermata, Tunicata and *Drosophila melanogaster* non-redundant protein database (using NCBI blast tool). Aligned sequences were retrieved using vertebrate genes names as an indication to cover putative ingroup and outgroups relative to the ascidians genes. Then alignments were performed using Mafft, trimmed and cleaned manually using BioEdit and analyzed using PhyML under the GTR+G+I model. Trees were visually rooted using treeview. In the cases where the retrieved sequences didn’t span over the outgroup, then more distant sequences were added to the alignment and the analysis were re-run. Then ascidian gene orthology was assigned by defining the smallest clade including the target *P. zorritensis* and *C. lepadiformis* contigs together with vertebrate sequences. All the alignements, the resulting phylogenies and the sequence accession numbers are provided in Supplementary Figure 3.

## Acknowledgements

We would like to thank the “Service Aquariologie” supported by EMBRC-France for the help with the marine-culture facility. We thank Marie Borriglione for her help with the environmental stress experiments, Philippe Dru for help with the bioinformatics resources and Sonia Lotito for the technical support. This study was supported by Sorbonne University EMERGENCE 2021/22 and INSB-DBM2021 to S.T.; FAPESP BEPE (2018/05923-3) to L. S. H.; FAPESP JP 2015/50164-5, 2018/50017-0 and CNPq 306638/2020-7 to F. D. B.

---

**Supplementary Figure 1:**
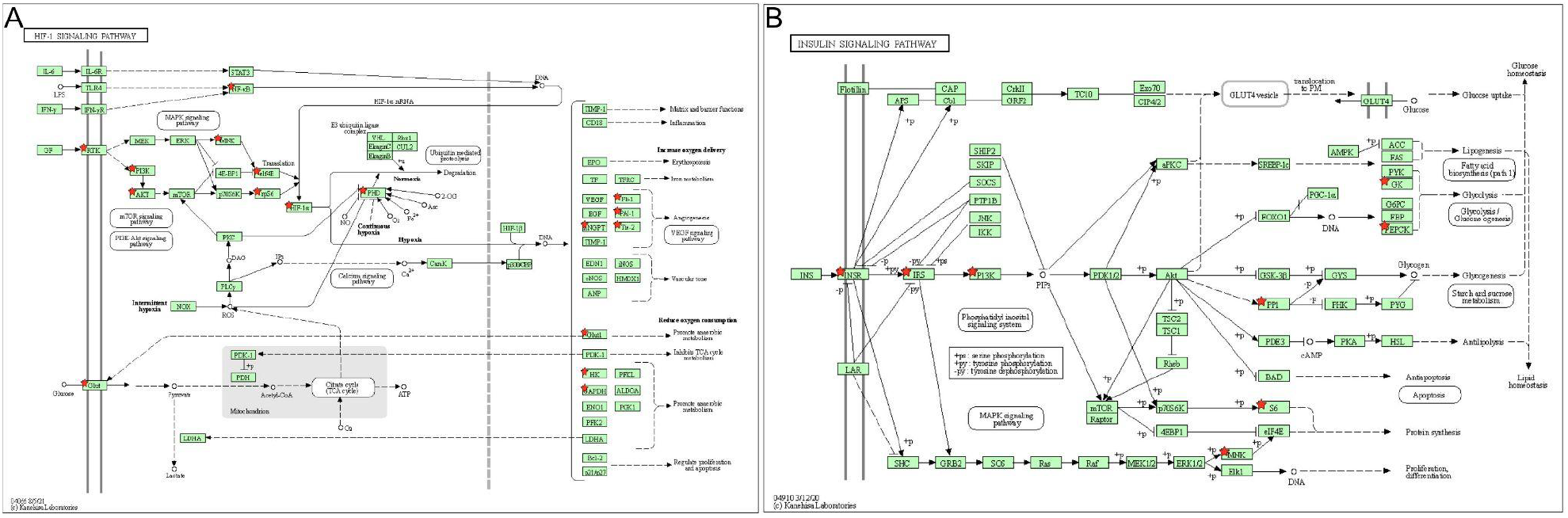
KEGG pathway diagrams for selected pathways: (A) HIF-1α signaling pathway and (B) insulin signaling pathway; red stars indicate genes that are upregulated in both species during dormancy.

**Supplementary Figure 2:**
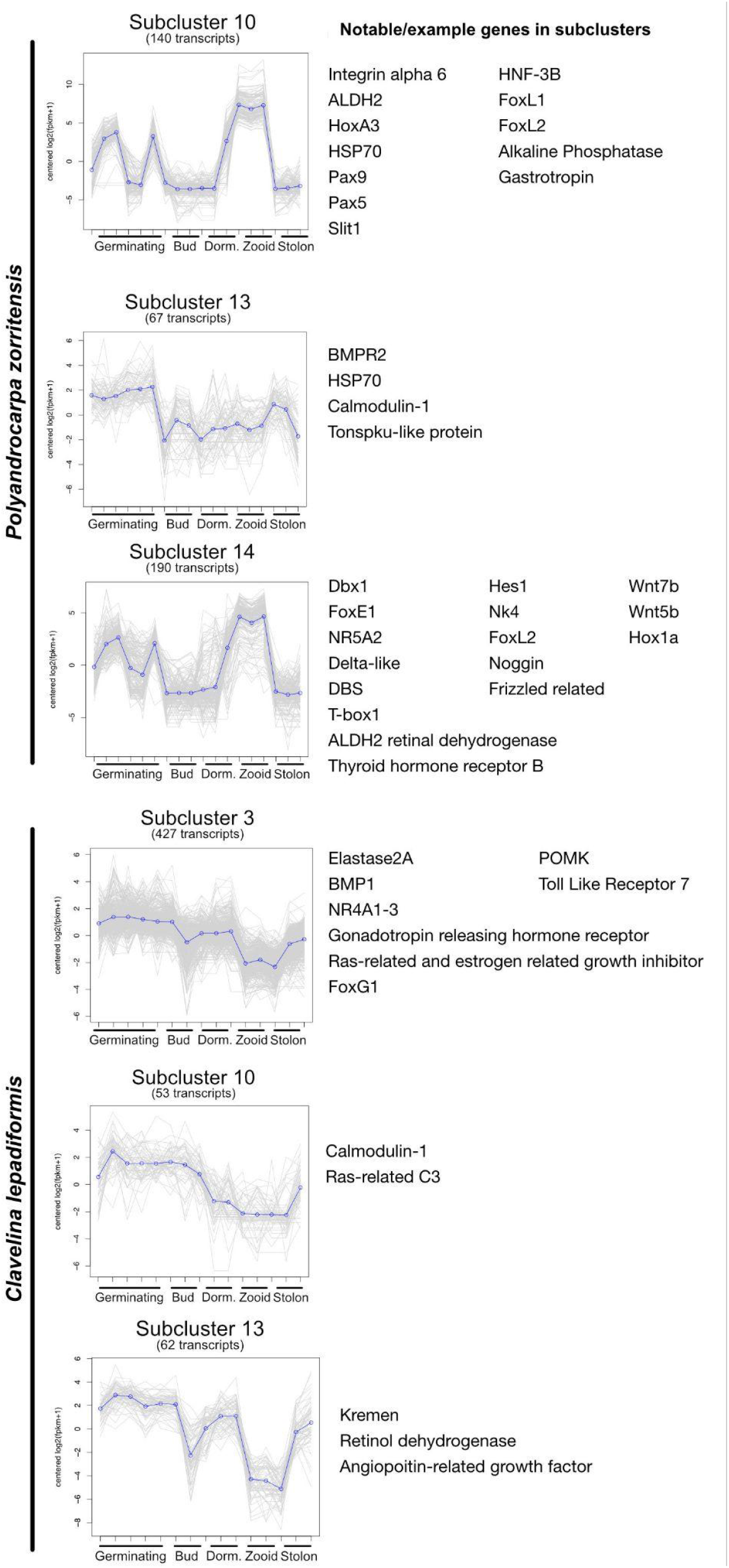
Highly expressed genes during germination in Polyandrocarpa zorritensis (top three subclusters) and Clavelina lepadiformis (bottom three subclusters). Representative genes worthy of mention for each subcluster (see Figs. 10-11) are shown on the right. Each gene is plotted (gray lines). The mean expression profile for that cluster is shown in blue. Expression values are shown as log2-transformed, median centered fpkm values. Subclusters were generated by cutting the hierarchical expression cluster tree by 30% tree height. Number of transcripts within each subcluster is indicated. Life stages are indicated on x-axis of each cluster: “Bud” is short for the “prebud” stage (budding chamber); “Dorm” is short for the dormant state (the winter bud).

