## Supplementary Figure 3 for "Comparing dormancy in two distantly related tunicates reveals morphological, molecular, and ecological convergences and repeated co-option"

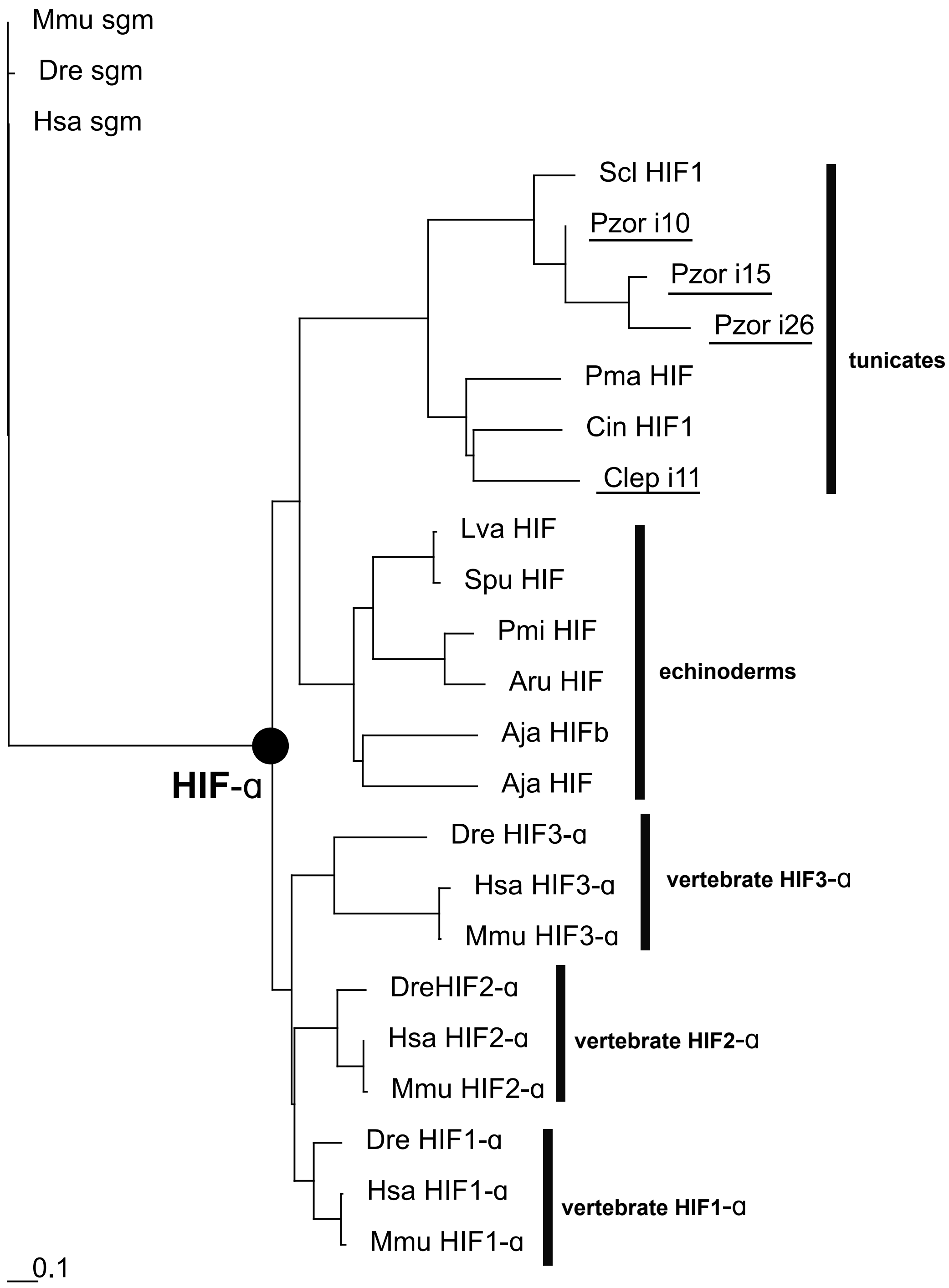
Mmu sgm
Dre sgm
Hsa sgm
Scl HIF1
Pzor i10
Pzor i15
Pzor i26
tunicates
Pma HIF
Cin HIF1
Clep i11
Lva HIF
Spu HIF
Pmi HIF
Aru HIF
echinoderms
Aja HIFb
Aja HIF
HIF-α
Dre HIF3-α
Hsa HIF3-α
Mmu HIF3-α
vertebrate HIF3-α
DreHIF2-α
Hsa HIF2-α
Mmu HIF2-α
vertebrate HIF2-α
Dre HIF1-α
Hsa HIF1-α
Mmu HIF1-α
vertebrate HIF1-α
0.1

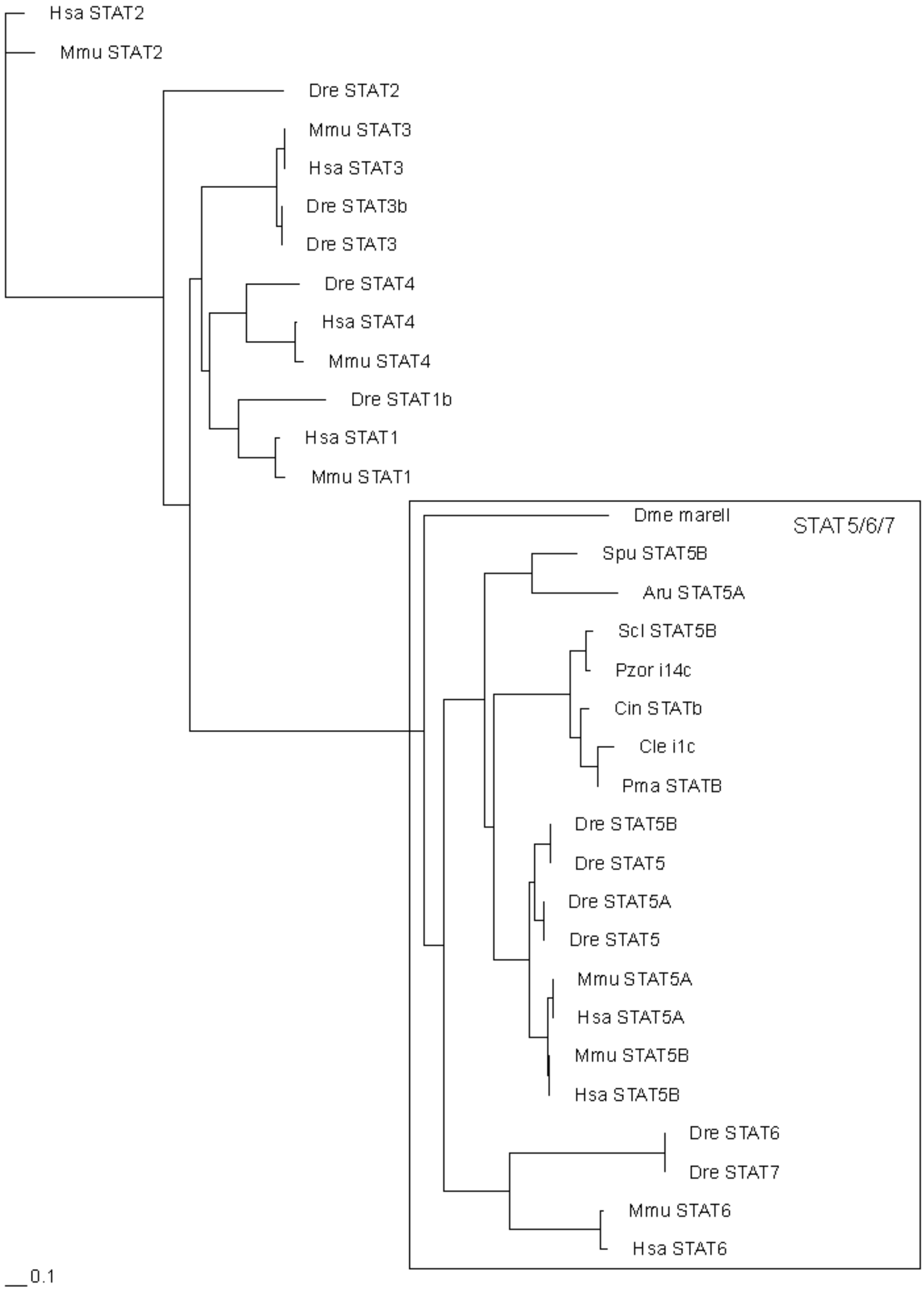
Hsa STAT2
Mmu STAT2
Dre STAT2
Mmu STAT3
Hsa STAT3
Dre STAT3b
Dre STAT3
Dre STAT4
Hsa STAT4
Mmu STAT4
Dre STAT1b
Hsa STAT1
Mmu STAT1
Dme marell
STAT5/6/7
Spu STAT5B
Aru STAT5A
Scl STAT5B
Pzor i14c
Cin STATb
Cle i1c
Pma STATB
Dre STAT5B
Dre STAT5
Dre STAT5A
Dre STAT5
Mmu STAT5A
Hsa STAT5A
Mmu STAT5B
Hsa STAT5B
Dre STAT6
Dre STAT7
Mmu STAT6
Hsa STAT6
0.1

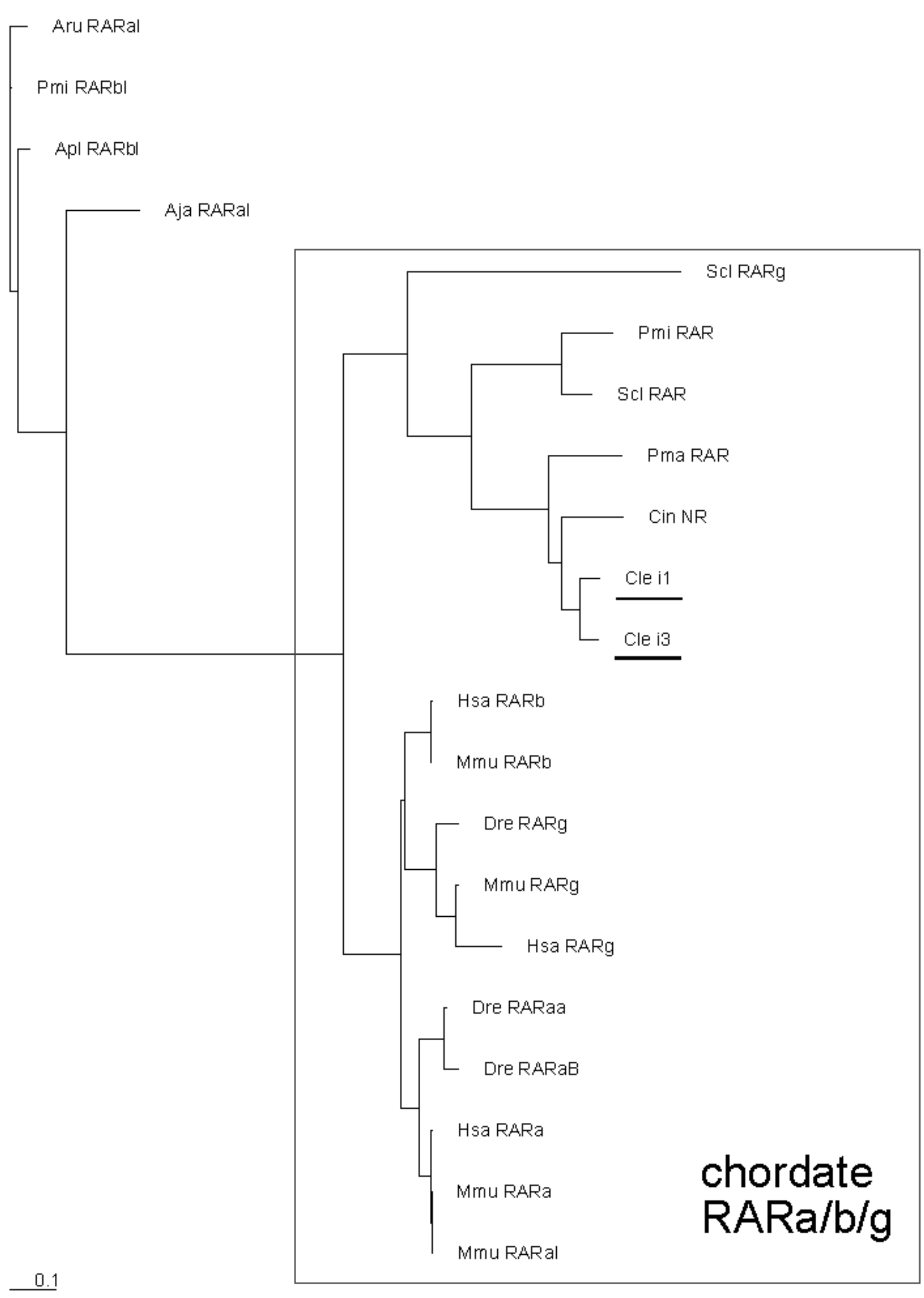
Aru RARal
Pmi RARbl
Apl RARbl
Aja RARal
Scl RARg
Pmi RAR
Scl RAR
Pma RAR
Cin NR
Cle i1
Cle i3
Hsa RARb
Mmu RARb
Dre RARg
Mmu RARg
Hsa RARg
Dre RARaa
Dre RARaB
Hsa RARa
Mmu RARa
Mmu RARal
chordate RARa/b/g
0.1

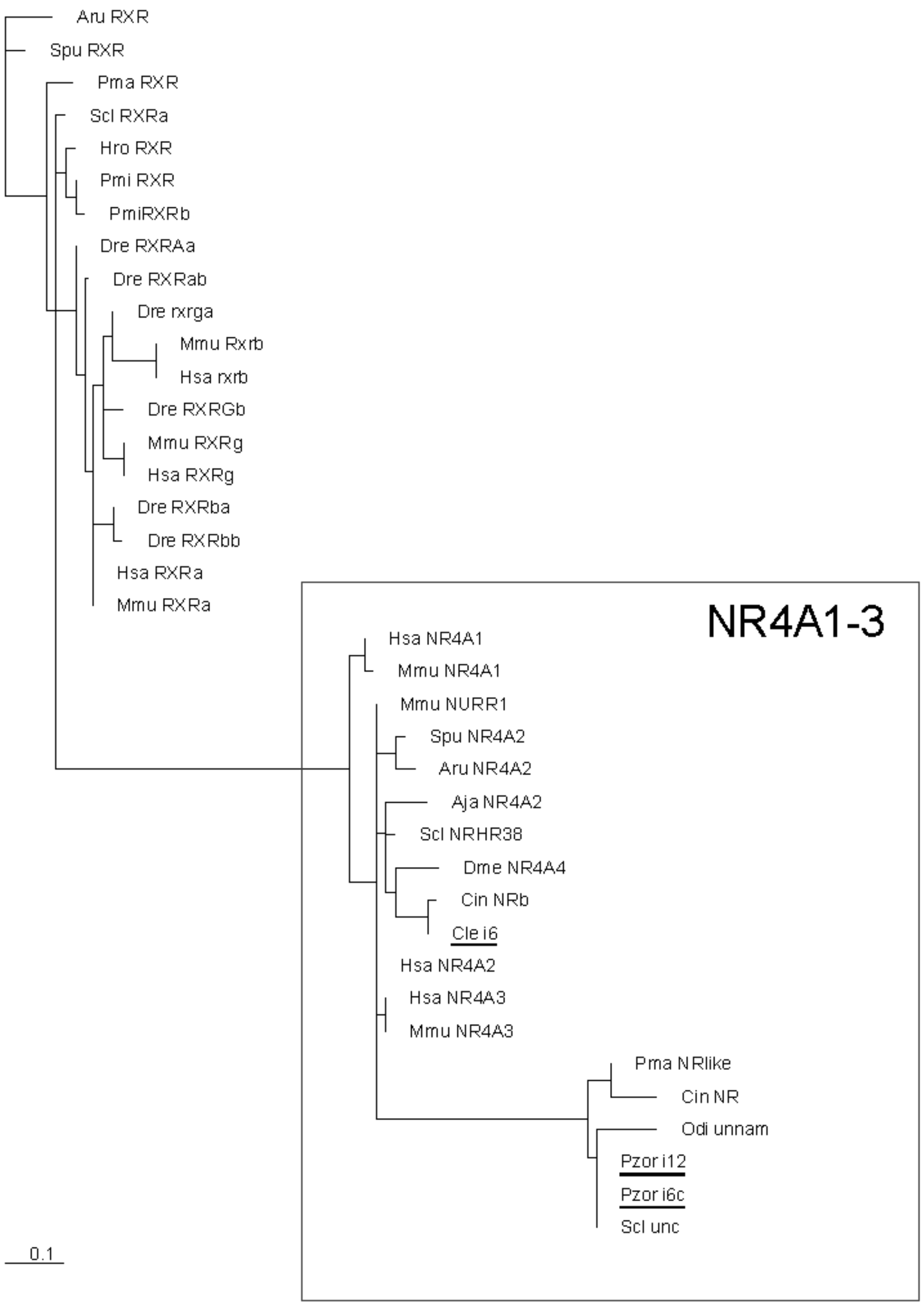

Aru RXR
Spu RXR
Pma RXR
Scl RXRa
Hro RXR
Pmi RXR
PmiRXRb
Dre RXRAa
Dre RXRab
Dre rxrga
Mmu Rxrb
Hsa rxrb
Dre RXRGb
Mmu RXRg
Hsa RXRg
Dre RXRba
Dre RXRbb
Hsa RXRa
Mmu RXRa
NR4A1-3
Hsa NR4A1
Mmu NR4A1
Mmu NURR1
Spu NR4A2
Aru NR4A2
Aja NR4A2
Scl NRHR38
Dme NR4A4
Cin NRb
Cle i6
Hsa NR4A2
Hsa NR4A3
Mmu NR4A3
Pma NRlike
Cin NR
Odi unnam
Pzor i12
Pzor i6c
Scl unc
0.1

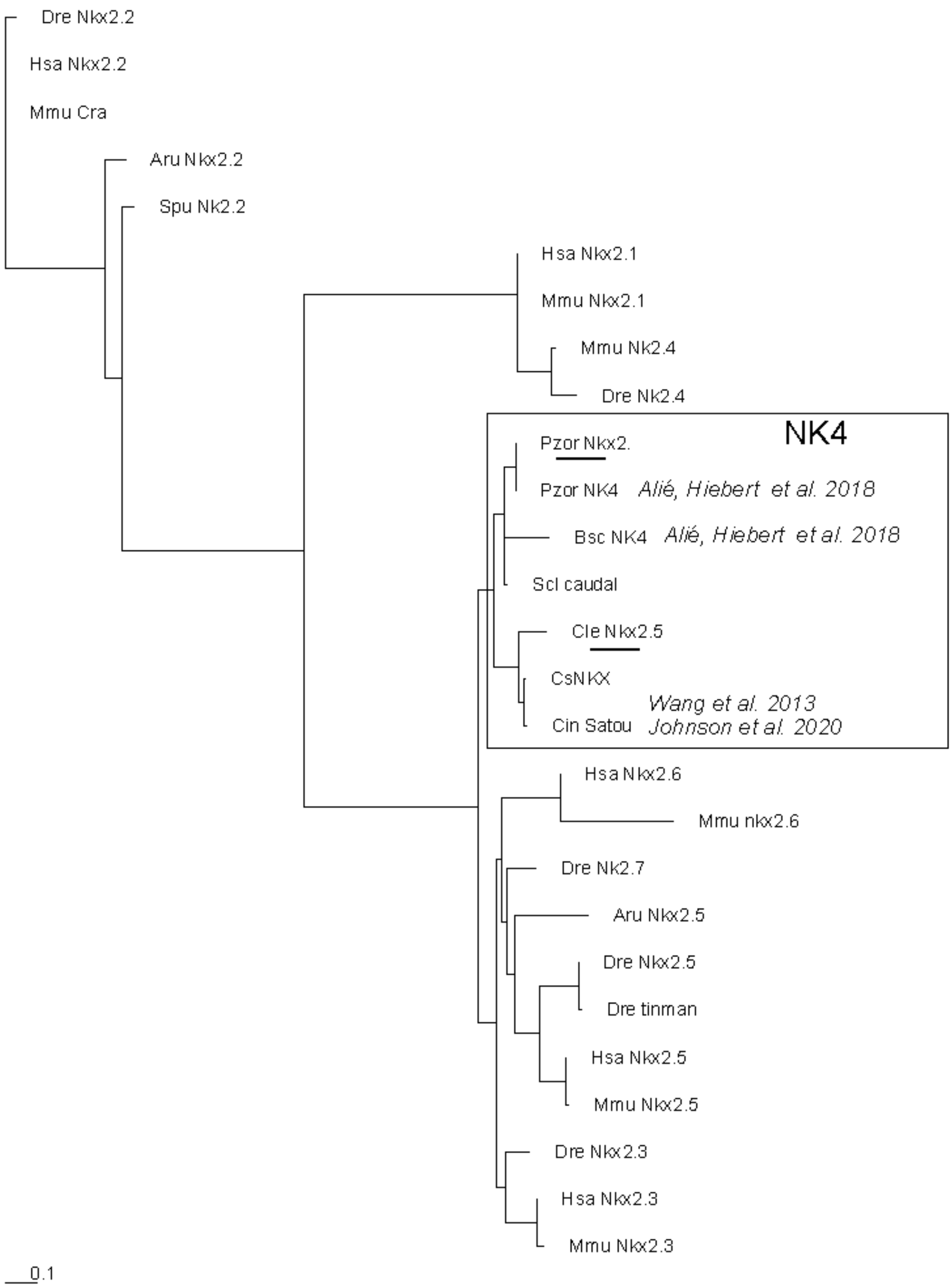

Dre Nkx2.2
Hsa Nkx2.2
Mmu Cra
Aru Nkx2.2
Spu Nk2.2
Hsa Nkx2.1
Mmu Nkx2.1
Mmu Nk2.4
Dre Nk2.4
NK4
Pzor Nkx2.
Pzor NK4
Alié, Hiebert et al. 2018
Bsc NK4
Alié, Hiebert et al. 2018
Scl caudal
Cle Nkx2.5
CsNKX
Wang et al. 2013
Cin Satou
Johnson et al. 2020
Hsa Nkx2.6
Mmu nkx2.6
Dre Nk2.7
Aru Nkx2.5
Dre Nkx2.5
Dre tinman
Hsa Nkx2.5
Mmu Nkx2.5
Dre Nkx2.3
Hsa Nkx2.3
Mmu Nkx2.3
0.1

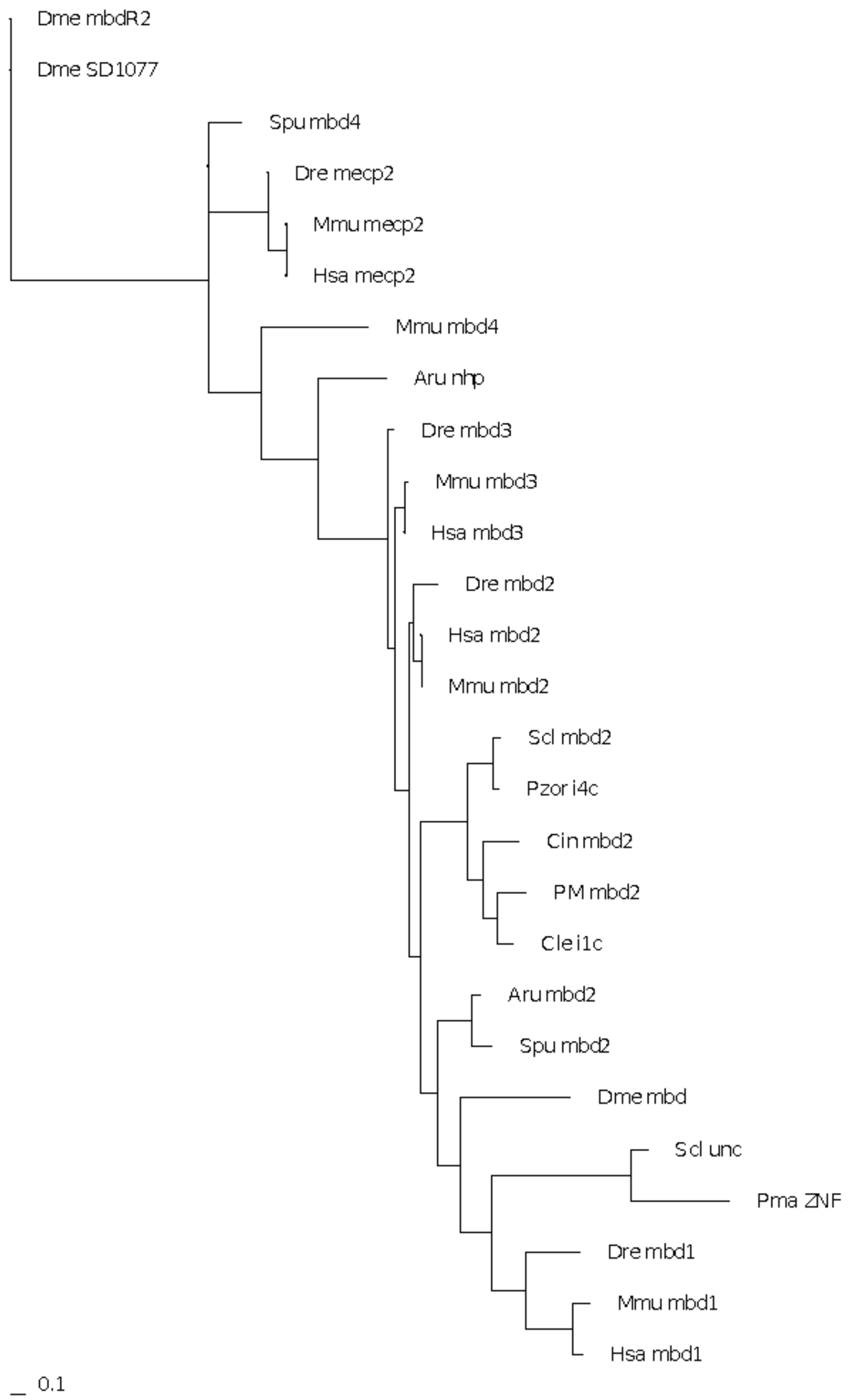
Dme mbdR2
Dme SD1077
Spu mbd4
Dre mecp2
Mmu mecp2
Hsa mecp2
Mmu mbd4
Aru nhp
Dre mbd3
Mmu mbd3
Hsa mbd3
Dre mbd2
Hsa mbd2
Mmu mbd2
Scl mbd2
Pzor i4c
Cin mbd2
PM mbd2
Cle i1c
Aru mbd2
Spu mbd2
Dme mbd
Scl unc
Pma ZNF
Dre mbd1
Mmu mbd1
Hsa mbd1
0.1

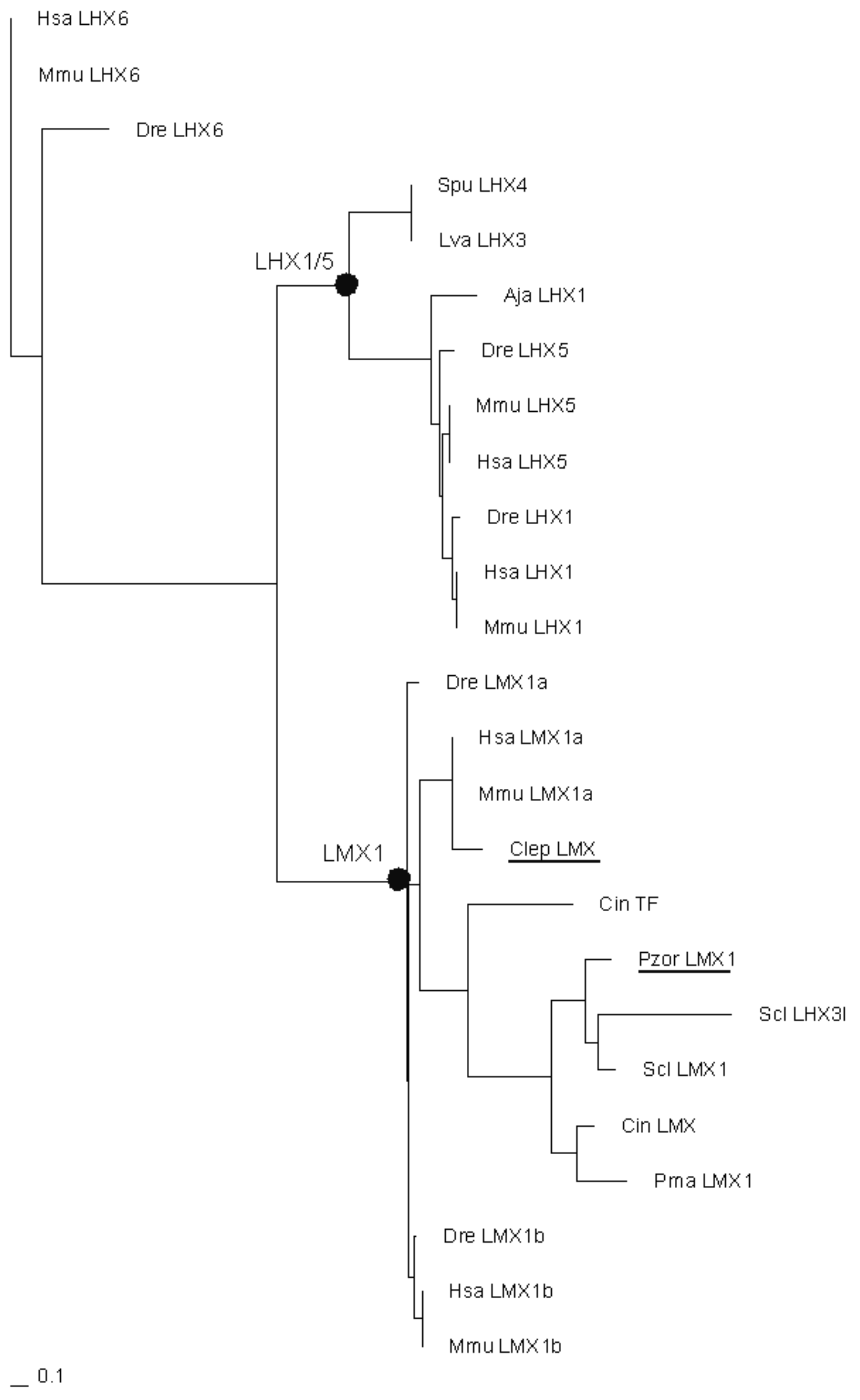
Hsa LHX6
Mmu LHX6
Dre LHX6
Spu LHX4
Lva LHX3
LHX1/5
Aja LHX1
Dre LHX5
Mmu LHX5
Hsa LHX5
Dre LHX1
Hsa LHX1
Mmu LHX1
Dre LMX1a
Hsa LMX1a
Mmu LMX1a
Clep LMX
LMX1
Cin TF
Pzor LMX1
Scl LHX3l
Scl LMX1
Cin LMX
Pma LMX1
Dre LMX1b
Hsa LMX1b
Mmu LMX1b
0.1

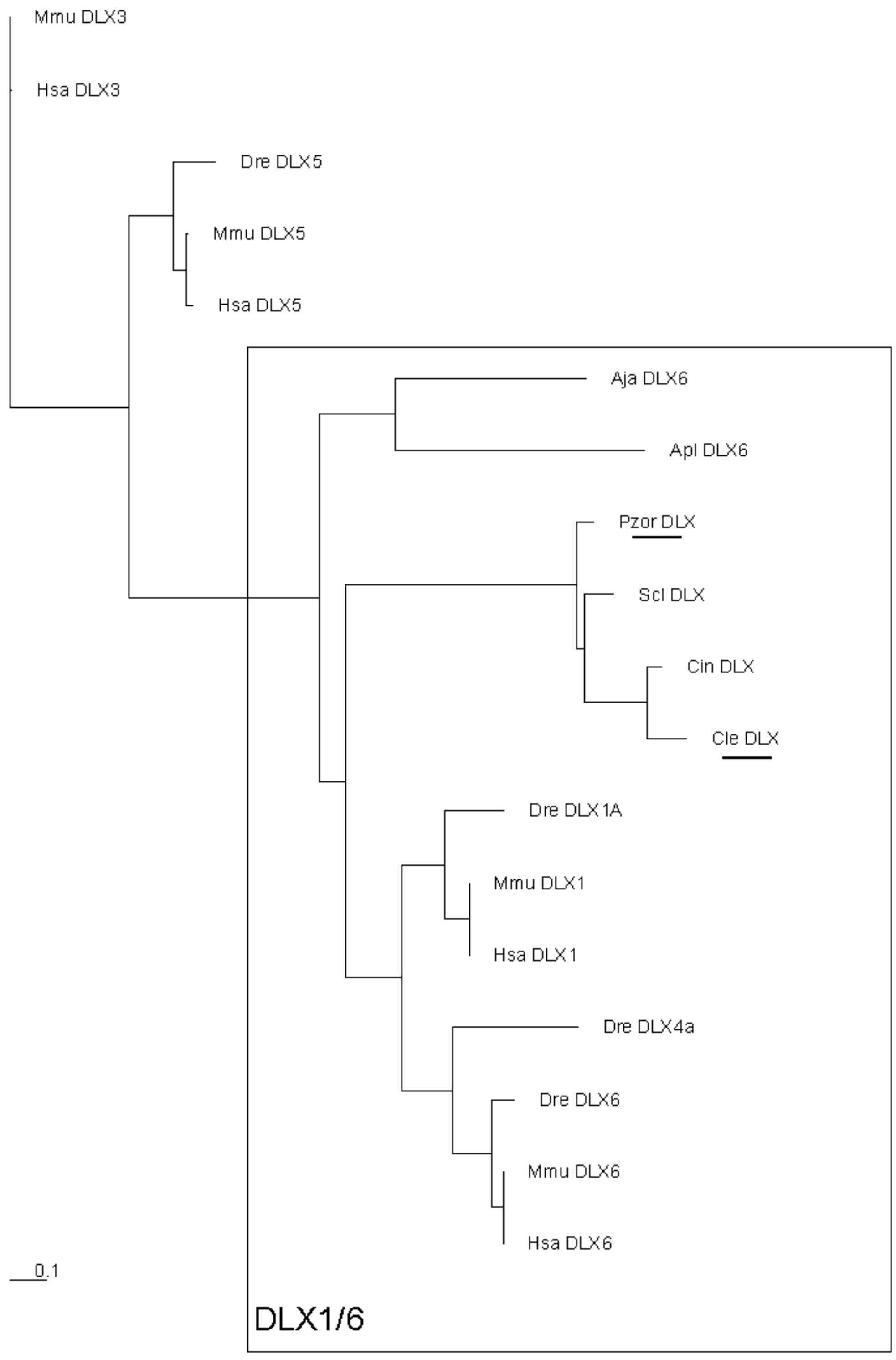
Mmu DLX3
Hsa DLX3
Dre DLX5
Mmu DLX5
Hsa DLX5
Aja DLX6
Apl DLX6
Pzor DLX
Scl DLX
Cin DLX
Cle DLX
Dre DLX1A
Mmu DLX1
Hsa DLX1
Dre DLX4a
Dre DLX6
Mmu DLX6
Hsa DLX6
0.1
DLX1/6

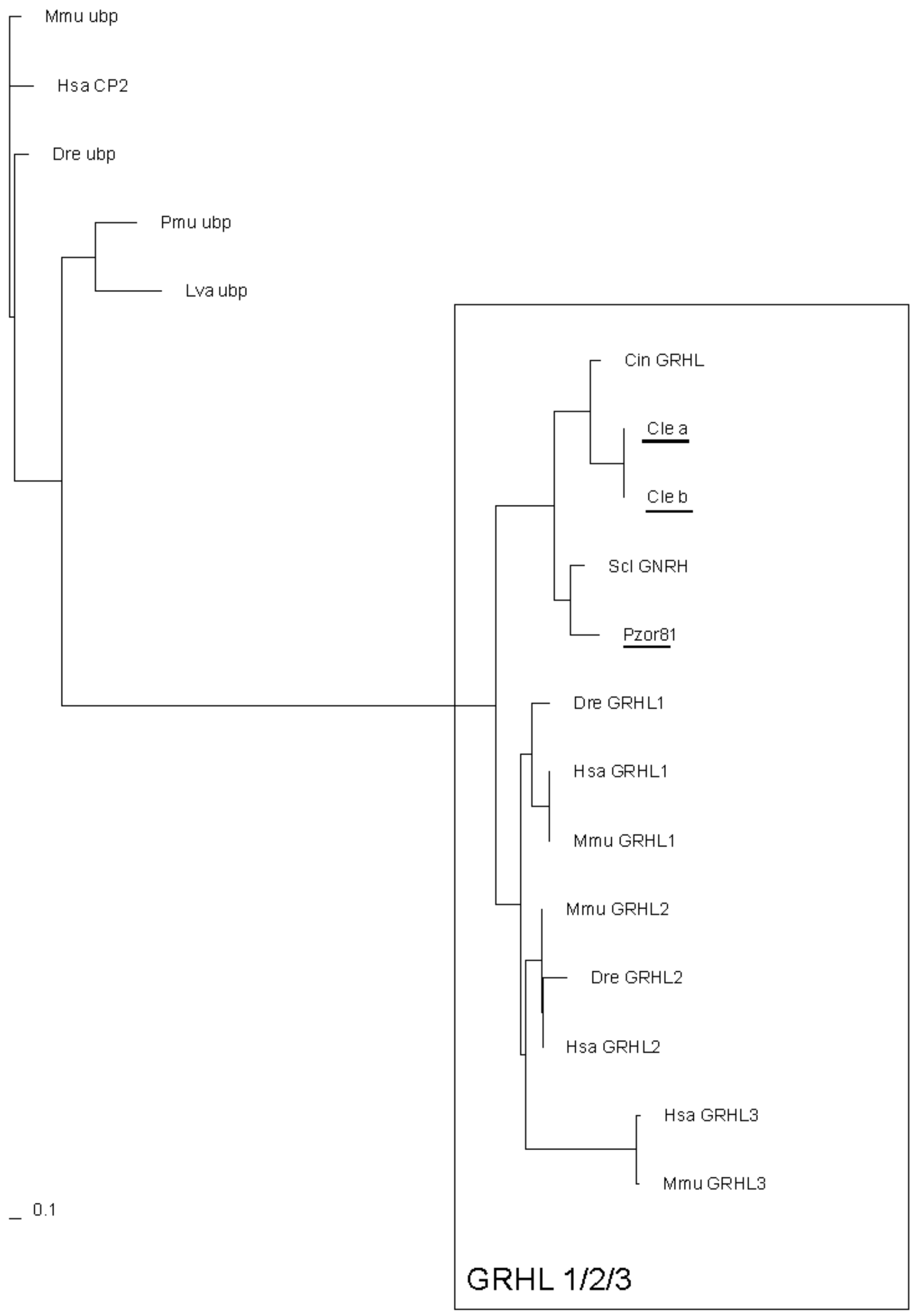

Mmu ubp
Hsa CP2
Dre ubp
Pmu ubp
Lva ubp
Cin GRHL
Cle a
Cle b
Scl GNRH
Pzor81
Dre GRHL1
Hsa GRHL1
Mmu GRHL1
Mmu GRHL2
Dre GRHL2
Hsa GRHL2
Hsa GRHL3
Mmu GRHL3
0.1
GRHL 1/2/3

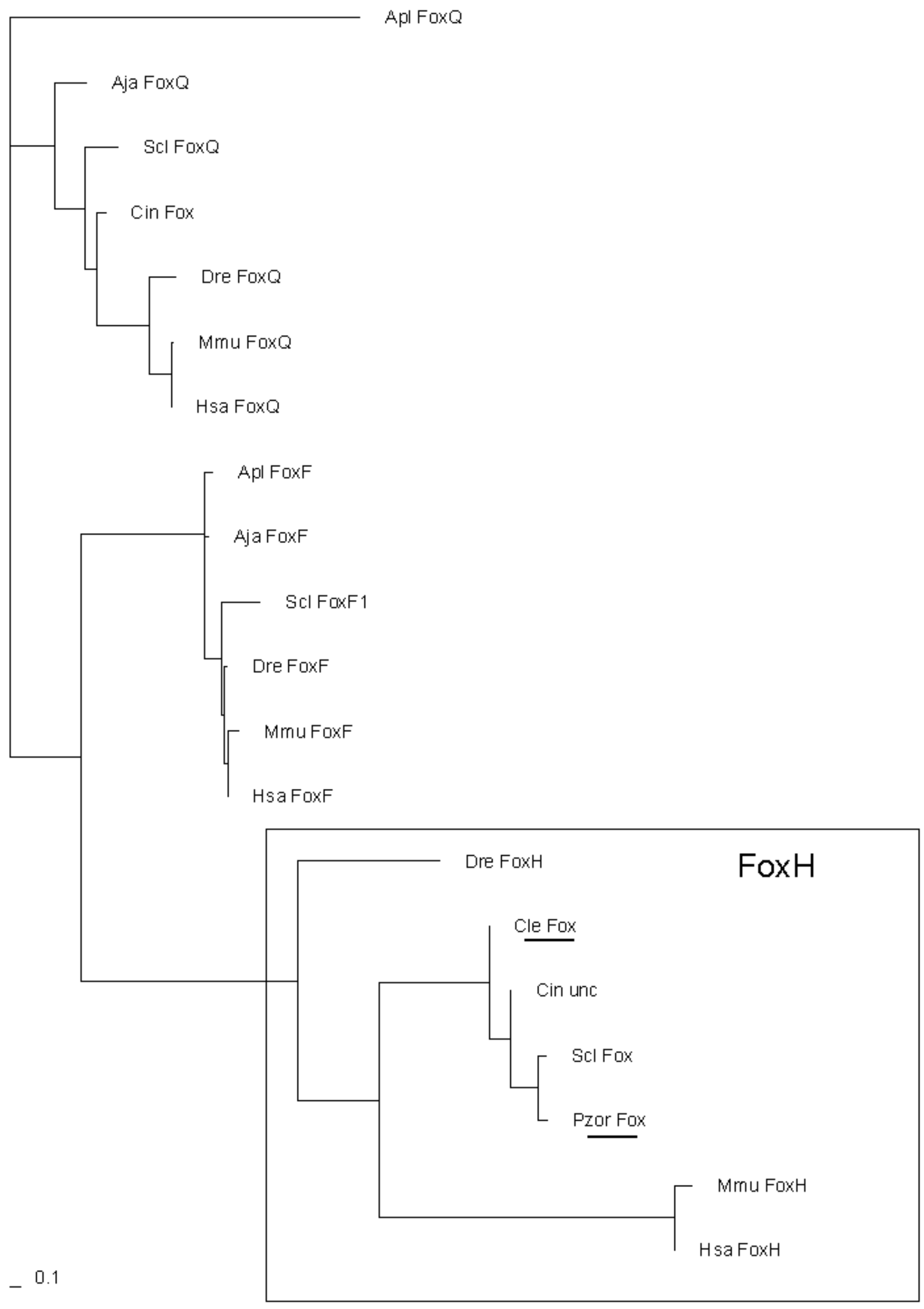
Apl FoxQ
Aja FoxQ
Scl FoxQ
Cin Fox
Dre FoxQ
Mmu FoxQ
Hsa FoxQ
Apl FoxF
Aja FoxF
Scl FoxF1
Dre FoxF
Mmu FoxF
Hsa FoxF
Dre FoxH
FoxH
Cle Fox
Cin unc
Scl Fox
Pzor Fox
Mmu FoxH
Hsa FoxH
0.1

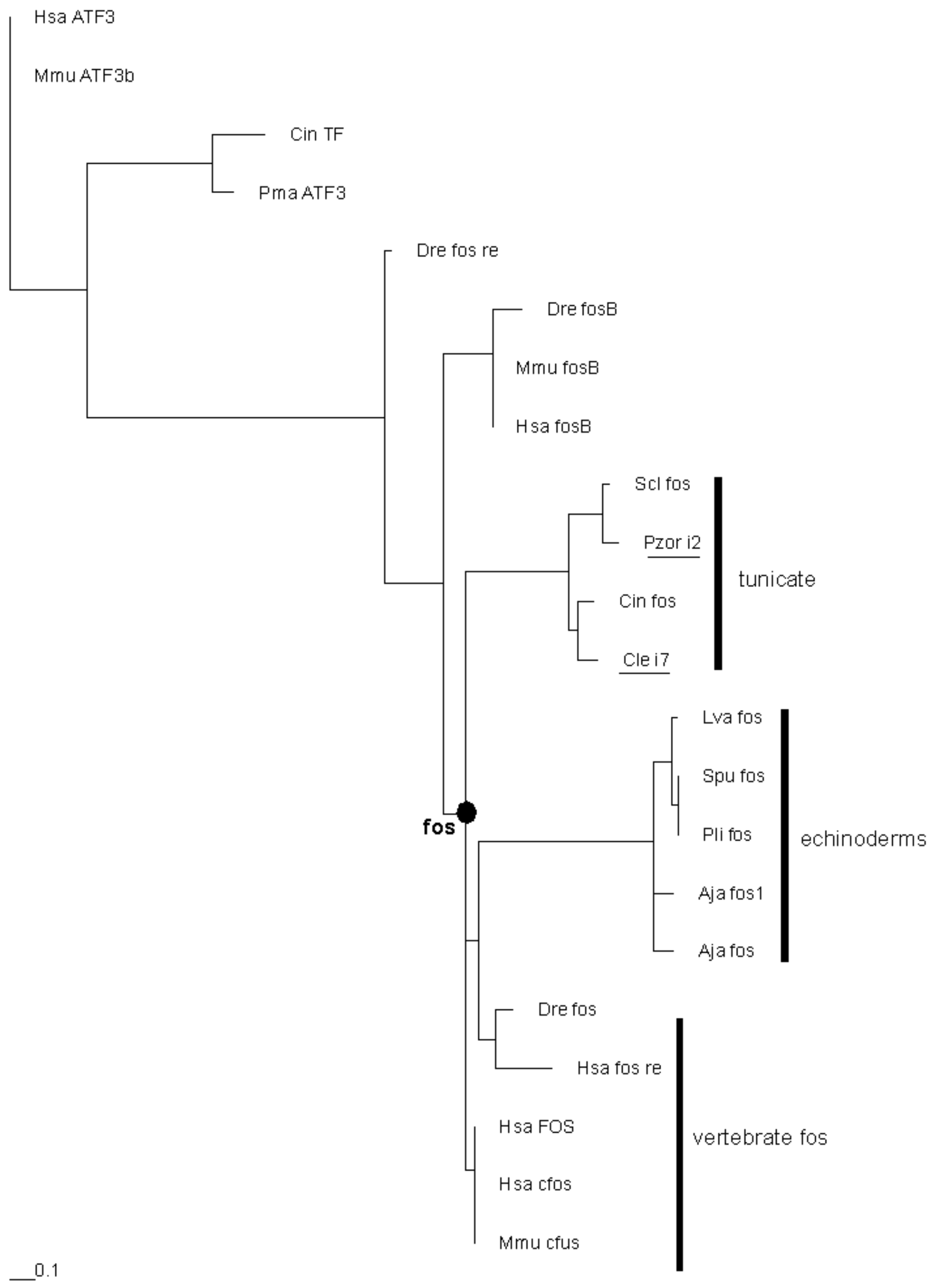
Hsa ATF3
Mmu ATF3b
Cin TF
Pma ATF3
Dre fos re
Dre fosB
Mmu fosB
Hsa fosB
Scl fos
Pzor i2
Cin fos
Cle i7
tunicate
Lva fos
Spu fos
Pli fos
Aja fos1
Aja fos
echinoderms
fos
Dre fos
Hsa fos re
Hsa FOS
Hsa cfos
Mmu cfus
vertebrate fos
0.1

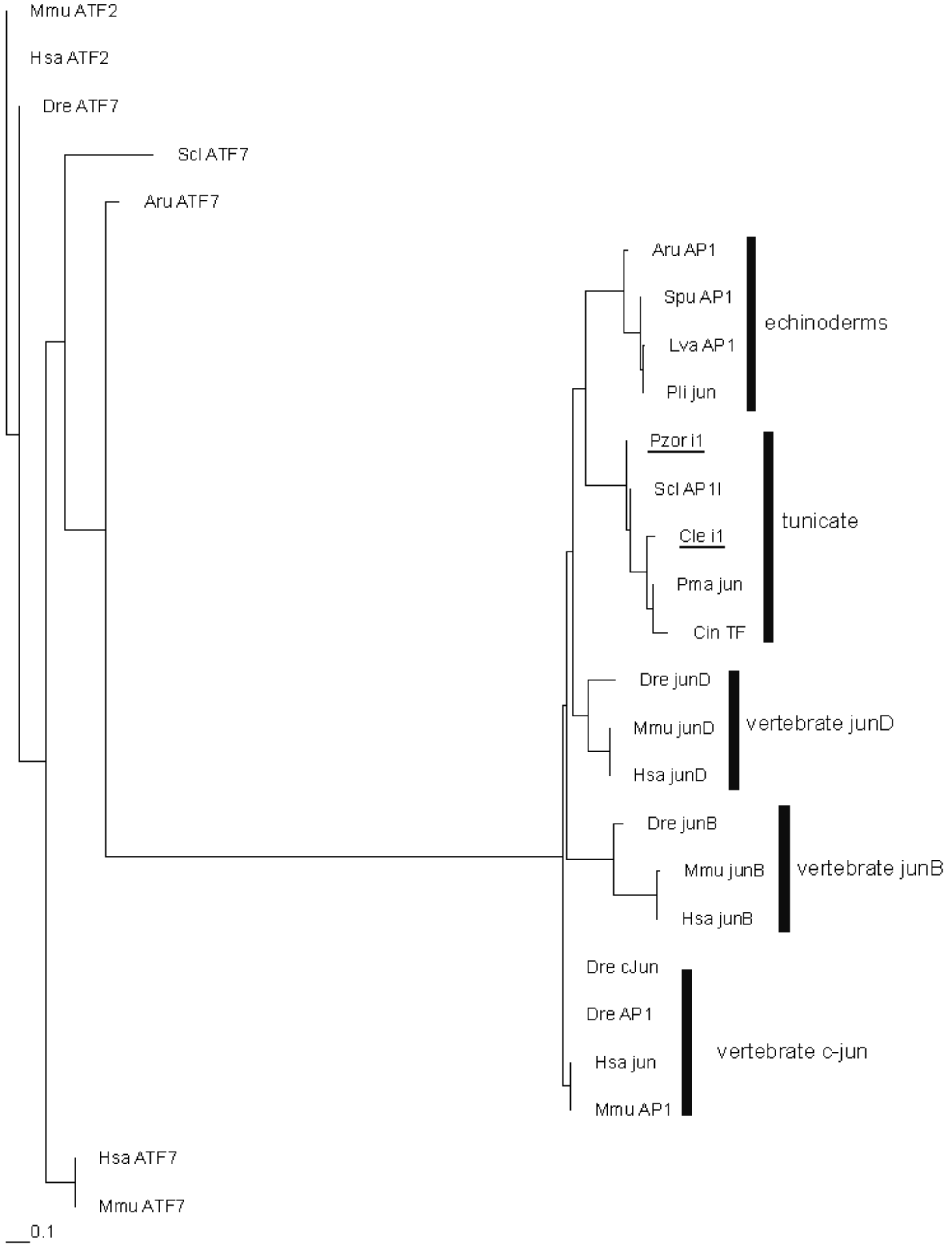
Mmu ATF2
Hsa ATF2
Dre ATF7
Scl ATF7
Aru ATF7
Aru AP1
Spu AP1
Lva AP1
Pli jun
echinoderms
Pzor i1
Scl AP1l
Cle i1
Pma jun
Cin TF
tunicate
Dre junD
Mmu junD
Hsa junD
vertebrate junD
Dre junB
Mmu junB
Hsa junB
vertebrate junB
Dre cJun
Dre AP1
Hsa jun
Mmu AP1
vertebrate c-jun
Hsa ATF7
Mmu ATF7
0.1

Abbreviations:

| | |
|---|---|
| Hsa | *Homo sapiens* |
| Mmu | *Mus musculus* |
| Dre | *Danio rerio* |
| Pzor | *Polyandrocarpa zorritensis* |
| Cle | *Clavelina lepadiformis* |
| Scl | *Styela clava* |
| Pma | *Phallusia mammillata* |
| Cin | *Ciona intestinalis* |
| Aru | *Asterias rubens* |
| Spu | *Strongylocentrotus purpuratus* |
| Lva | *Lytechinus variegatus* |
| Ana | *Anneissia japonica* |
| Pmi | *Patiria miniata* |
| Dme | *Drosophila melanogaster* |
| Apl | *Acanthaster planci* |
| Pli | *Paracentrotus lividus* |
| Odi | *Oikopleura dioica* |
| Hro | *Halocynthia roretzi* |
| Pmi | *Polyandrocarpa misakiensis* |

| Name in the tree | Accession |
|---|---|
| HIF tree | |
| Scl_HIF1 | XP_039247709.1 |
| Pzor_i10 | Pzorritensis_TRINITY_DN1515_c0_g1_i10c |
| Pzor_i15 | Pzorritensis_TRINITY_DN1515_c0_g1_i15c |
| Pzor_i26 | Pzorritensis_TRINITY_DN1515_c0_g1_i26c |
| Cin_HIF1 | NP_001071731.1 |
| Pma_HIF | CAB3252826.1 |
| Dre_HIF1 | NP_001296971.1 |
| Hsa_HIF1 | NP_851397.1 |
| Mmu_HIF1 | AAC52730.1 |
| Hsa_PAS | NP_001421.2 |
| Mmu_PAS | AAH57870.1 |
| Dre_PAS | NP_001034895.2 |
| Dre_HIF3 | XP_005157629.2 |
| Hsa_HIF3 | XP_005259212.1 |
| Mmu_HIF3 | AAF21782.1 |
| Lva_HIF | XP_041468476.1 |
| Spu_HIF | XP_030854578.1 |
| Aja_HIF | QOE88730.1 |

| | |
|---|---|
| Pmi_HIF | XP_038074638.1 |
| Aru_HIF | XP_033631701.1 |
| Aja_HIF | XP_033104914.1 |
| Clep_i11 | Clepadiformis_TRINITY_DN6721_c0_g2_i11c |
| Hsa_sgm | XP_011534374.1 |
| Mmu_sgm | NP_035506.2 |
| Dre_sgm | NP_835740.2 |
| HLF tree | |
| Scl_HLF | XP_039256594.1 |
| Cin_TF | NP_001071833.1 |
| Pma_HLF | CAB3253308.1 |
| Cle_i14c | Clepadiformis_TRINITY_DN3733_c4_g1_i14c |
| Pzor_i5c | Pzorritensis_TRINITY_DN368_c1_g1_i5c |
| Pzor_i6c | Pzorritensis_TRINITY_DN368_c1_g1_i6c |
| Mmu_HLF | EDL15886.1 |
| Hsa_HLF | NP_002117.1 |
| Dre_TEF | NP_571475.1 |
| Mmu_TEF | XP_030104321.1 |
| Hsa_TEF | AAA81373.1 |
| Dre_HLF | NP_001183988.1 |
| Dre_Dsb | XP_005157954.1 |
| Dre_HLF | NP_001070802.1 |
| Mmu_Dsb | XP_006540659.1 |
| Hsa_Dsb | XP_016881877.1 |
| Pmi_TEF | XP_038055420.1 |
| Apl_TEF | XP_022103079.1 |
| Aru_TEF | XP_033631572.1 |
| Lva_TEF | XP_041454174.1 |
| Lhx1 tree | |
| Scl_LMX1 | XP_039269611.1 |
| Pzor_LMX1 | Pzorritensis_TRINITY_DN5540_c0_g1_i2c |
| Clep_LMX | Clepadiformis_TRINITY_DN10436_c0_g1_i1c |
| Cin_LMX | NP_001071998.1 |
| Pma_LMX1 | CAB3263460.1 |
| Hsa_LMX1a | XP_011507842.1 |
| Mmu_LMX1a | NP_387501.1 |
| Dre_LMX1a | NP_001020339.1 |
| Hsa_LMX1b | AAC39738.1 |
| Mmu_LMX1b | NP_034855.3 |
| Dre_LMX1b | NP_001020338.2 |
| Scl_LHX3l | XP_039270412.1 |
| Cin_TF | NP_001071756.1 |

| | |
|---|---|
| Hsa_LHX1 | AAH20470.1 |
| Mmu_LHX1 | NP_032524.1 |
| Dre_LHX1 | NP_571291.1 |
| Mmu_LHX5 | NP_032525.1 |
| Hsa_LHX5 | NP_071758.1 |
| Dre_LHX5 | NP_571293.1 |
| Aja_LHX1 | XP_033123538.1 |
| Spu_LHX4 | XP_030852869.1 |
| Lva_LHX3 | XP_041481746.1 |
| Dre_LHX6 | XP_685757.5 |
| Hsa_LHX6 | NP_001335119.1 |
| Mmu_LHX6 | NP_001076596.1 |
| fos tree | |
| Scl_fos | XP_039253751.1:48-562 |
| Cin_fos | XP_009860203.2:21-424 |
| Hsa_cfos | NP_005243.1:71-209 |
| Hsa_FOS | AKR52853.1:1-42 |
| Mmu_cfus | NP_034364.1:71-209 |
| Dre_fos | NP_001155024.1:125-202 |
| Mmu_fosB | NP_001334515.1:170-224 |
| Hsa_fosB | XP_005258748.1:170-224 |
| Dre_fosB | NP_001315131.1:213-282 |
| Hsa_fos_re | NP_001287785.1:54-138 |
| Dre_fos_re | NP_001349536.1:20-155 |
| Pzor_i2 | fos_Pzorritensis_TRINITY_DN2975_c0_g1_i2 |
| Aja_fos | XP_033104004.1:114-193 |
| Lva_fos | XP_041468790.1:144-222 |
| Spu_fos | XP_011679928.1:114-188 |
| Pli_fos | CZR14360.2:173-226 |
| Aja_fos | PIK60907.1:26-119 |
| Cle_i7 | fos_Clepadiformis_TRINITY_DN3286_c0_g1_i7c |
| Hsa_ATF3 | NP_001193413.2:44-98 |
| Mmu_ATF3b | AAP92420.1:44-98 |
| Cin_TF | NP_001071658.1:159-242 |
| Pma_ATF3 | CAB3224128.1:153-207 |
| jun tree | |
| Scl_AP1l | XP_039273144.1:1-381 |
| Pzor_i1 | Pzorritensis_TRINITY_DN9568_c0_g2_i1c |
| Pma_jun | CAB3257837.1:1-417 |
| Cin_TF | NP_001071996.1:1-381 |
| Dre_AP1 | NP_956281.1:57-308 |
| Dre_cJun | AAY21257.1:57-308 |

| | |
|---|---|
| Hsa_jun | AAH09874.2:120-231 |
| Mmu_AP1 | NP_034721.1:223-334 |
| Mmu_junD | NP_001273873.1:182-293 |
| Hsa_junD | NP_001273897.1:193-304 |
| Dre_junD | NP_001121814.1:176-282 |
| Dre_junB | ACR83585.1:183-301 |
| Mmu_junB | AAA74916.1:234-344 |
| Hsa_junB | AAH09465.1:237-347 |
| Lva_AP1 | XP_041474997.1:41-309 |
| Pli_jun | AIF71194.1:42-307 |
| Spu_AP1 | XP_030838976.1:43-309 |
| Aru_AP1 | XP_033637193.1:38-321 |
| Cle_i1 | Clepadiformis_TRINITY_DN6146_c0_g1_i1c |
| Hsa_ATF7 | NP_001353485.1:336-395 |
| Mmu_ATF7 | NP_001296999.1:334-395 |
| Dre_ATF7 | XP_009302699.1:340-416 |
| Mmu_ATF2 | NP_001271305.1:288-364 |
| Hsa_ATF2 | AAY17208.1:247-323 |
| Aru_ATF7 | XP_033624955.1:329-406 |
| Scl_ATF7 | XP_039266799.1:522-602 |
| Dre_BACH1 | NP_001035403.1:21-88 |
| RAR tree | |
| Pma_RAR | CAB3265442.1 |
| Cin_NR | NP_001072037.1 |
| Cle_i1 | Clepadiformis_TRINITY_DN2210_c0_g2_i1c |
| Pmi_RAR | BAA13143.1 |
| Scl_RAR | XP_039274750.1 |
| Cle_i3 | Clepadiformis_TRINITY_DN2210_c0_g2_i3c |
| Dre_RARaa | NP_571481.2 |
| Dre_RARaB | NP_571474.1 |
| Hsa_RARa | XP_011523398.1 |
| Mmu_RARa | NP_001348883.1 |
| Hsa_RARb | AAH50415.2 |
| Mmu_RARb | NP_035373.1 |
| Mmu_RARal | CAA40749.1 |
| Mmu_RARg | AAA40035.1 |
| Dre_RARg | AAB59953.1 |
| Hsa_RARg | NP_001230661.1 |
| Scl_RARg | XP_039266672.1 |
| Aru_RARal | XP_033629029.1 |
| Apl_RARbl | XP_022109469.1 |
| Pmi_RARbl | XP_038071552.1 |

| | |
|---|---|
| Aja_RARal | XP_033104820.1 |
| NR4A tree | |
| Scl_unc | XP_039266535.1 |
| Odi_unnam | CBY38503.1 |
| Pma_NRlike | CAB3264439.1 |
| Pzor_i6c | Pzorritensis_TRINITY_DN512_c0_g1_i6c |
| Cin_NR | FAA00109.1 |
| Pzor_i12 | Pzorritensis_TRINITY_DN512_c0_g1_i12c |
| Scl_NRHR38 | XP_039268852.1 |
| Mmu_NURR1 | Q06219.1 |
| Hsa_NR4A2 | P43354.1 |
| Hsa_NR4A3 | Q92570.3 |
| Mmu_NR4A3 | Q9QZB6.1 |
| Spu_NR4A2 | XP_003725145.1 |
| Aru_NR4A2 | XP_033628939.1 |
| Hsa_NR4A1 | P22736.1 |
| Mmu_NR4A1 | P12813.1 |
| Cin_NRb | NP_001071779.1 |
| Cle_i6 | Clepadiformis_TRINITY_DN43_c0_g2_i6c |
| Aja_NR4A2 | XP_033127899.1 |
| Dme_NR4A4 | P49869.3 |
| Scl_RXRa | XP_039269445.1 |
| Dre_rxrga | Q90415.1 |
| Dre_RXRab | A2T929.2 |
| Hsa_RXRa | P19793.1 |
| Mmu_RXRa | P28700.1 |
| Dre_RXRAa | A2T929.2 |
| Dre_RXRba | Q7SYN5.1 |
| Dre_RXRbb | Q90417.1 |
| Pmi_RXR | BAA82618.1 |
| PmiRXRb | BAM66778.1 |
| Hro_RXR | AEW68002.1 |
| Mmu_RXRg | P28705.2 |
| Hsa_RXRg | P48443.1 |
| Dre_RXRGb | Q6DHP9.1 |
| Mmu_Rxrb | P28704.2 |
| Hsa_rxrb | P28702.2 |
| Spu_RXR | XP_030853250.1 |
| Aru_RXR | XP_033646607.1 |
| Pma_RXR | CAB3265841.1 |
| Stat tree | |
| Mmu_STAT5B | P42232.1 |

| | |
|---|---|
| Hsa_STAT5B | P51692.2 |
| Mmu_STAT5A | P42230.1 |
| Hsa_STAT5A | P42229.1 |
| Dre_STAT5A | NP_919368.2 |
| Dre_STAT5 | AAL73244.1 |
| Dre_STAT5B | NP_001003984.1 |
| Dre_STAT5_2 | AAI39518.1 |
| Scl_STAT5B | XP_039268530.1 |
| Pzor_i14c | Pzorritensis_TRINITY_DN4104_c0_g1_i14c |
| Cin_STATb | NP_001071828.1 |
| Pma_STATB | CAB3266641.1 |
| Cle_i1c | Clepadiformis_TRINITY_DN1191_c0_g2_i1c |
| Spu_STAT5B | XP_781834.3 |
| Aru_STAT5A | XP_033625377.1 |
| Mmu_STAT6 | P52633.2 |
| Hsa_STAT6 | P42226.1 |
| Dre_STAT6 | NP_001124062.1 |
| Dre_STAT7 | BAH47265.1 |
| Dme_marelle | Q24151.1 |
| Mmu_STAT3 | P42227.2 |
| Hsa_STAT3 | P40763.2 |
| Dre_STAT3b1 | AQT31669.1 |
| Dre_STAT3 | BAH47263.1 |
| Hsa_STAT1 | P42224.2 |
| Mmu_STAT1 | P42225.1 |
| Dre_STAT1b | NP_956385.2 |
| Hsa_STAT4 | Q14765.1 |
| Mmu_STAT4 | P42228.1 |
| Dre_STAT4 | XP_021334299.1 |
| Dre_STAT2 | NP_001258730.1 |
| Hsa_STAT2 | P52630.1 |
| Mmu_STAT2 | Q9WVL2.1 |
| MBD2 tree | |
| Scl_mbd2 | XP_039270644.1 |
| Pzor_i4c | Pzorritensis_TRINITY_DN24453_c0_g1_i4c |
| PM_mbd2 | CAB3263664.1 |
| Cle_i1c | Clepadiformis_TRINITY_DN10071_c0_g1_i1c |
| Cin_mbd2 | XP_002132141.1 |
| Dre_mbd2 | NP_997933.1 |
| Hsa_mbd2 | Q9UBB5.1 |
| Mmu_mbd2 | Q9Z2E1.2 |
| Dre_mbd3 | AAH71531.1 |

| | |
|---|---|
| Mmu_mbd3 | NP_997745.1 |
| Hsa_mbd3 | Q9D9H3.1 |
| Aru_mbd2 | XP_033633865.1 |
| Spu_mbd2 | XP_011678571.1 |
| Dme_mbd | NP_649907.1 |
| Mmu_mbd1 | Q9Z2E2.2 |
| Hsa_mbd1 | Q9UIS9.2 |
| Dre_mbd1 | AAI39876.1 |
| Aru_nhp | XP_033648028.1 |
| Dre_mecp2 | AAH93116.1 |
| Mmu_mecp2 | Q9Z2D6.1 |
| Hsa_mecp2 | P51608.1 |
| Spu_mbd4 | XP_030856411.1 |
| Mmu_mbd4 | Q9Z2D7.1 |
| Scl_unc | XP_039268150.1 |
| Pma_ZNF | CAB3263665.1 |
| Dme_mbdR2 | NP_731688.2 |
| Dme_SD10773 | AAM29652.1 |
| NK4 tree | |
| Scl_caudal | XP_039259202.1 |
| Dre_Nkx2.5 | AAH93132.1 |
| Dre_tinman | AAB50916.1 |
| Hsa_Nkx2.5 | NP_004378.1 |
| Mmu_Nkx2.5 | NP_032726.1 |
| Hsa_Nkx2.3 | NP_660328.2 |
| Mmu_Nkx2.3 | NP_032725.1 |
| Dre_Nkx2.3 | NP_571498.1 |
| Pzor_Nkx2.5 | NKX2-5_Pzorritensis_TRINITY_DN103704_c0_g1_i1c |
| Pzor_NK4 | Pzor_NK4_(Alié et al. 2018 - MBE) |
| Hsa_Nkx2.6 | NP_001129743.2 |
| Mmu_nkx2.6 | AAI00426.1 |
| Dre_Nk2.7 | AAH79493.1 |
| Cle_Nkx2.5 | NKX2-5_Clepadiformis_TRINITY_DN2235_c0_g1_i1c |
| CsNKX | BAA25399.1 |
| Cin_TF | NP_001071957.1_(Wang et al. 2013) |
| Bsc_NK4 | Bsc_NK4_(Alié et al 2018 - MBE) |
| Aru_Nkx2.5 | XP_033632513.1 |
| Aru_Nkx2.2 | XP_033640004.1 |
| Mmu_Cra | EDL28521.1 |
| Hsa_Nkx2.2 | NP_002500.1 |
| Dre_Nkx2.2 | NP_001295569.1 |
| Spu_Nk2.2 | AAS58444.1 |

| | |
|---|---|
| Dre_Nk2.4 | NP_571664.1 |
| Mmu_Nk2.4 | NP_075993.1 |
| Hsa_Nkx2.1 | NP_003308.1 |
| Mmu_Nkx2.1 | NP_033411.3 |
| GRHL tree | |
| Scl_GNRH | XP_039271768.1 |
| Pzor81 | Polyandrocarpa-GRHL1_TRINITY_DN81_c1_g1_i1.p1 |
| Cin_GRHL | XP_002131671.1 |
| Cle_a | Clavelina-GRHL1_TRINITY_DN5657_c0_g1_i5.p1a |
| Cle_b | Clavelina-GRHL1_TRINITY_DN5657_c0_g1_i5.p1c |
| Dre_GRHL1 | XP_017207086.1 |
| Hsa_GRHL1 | XP_011508645.1 |
| Mmu_GRHL1 | NP_001154878.1 |
| Mmu_GRHL2 | XP_006520126.3 |
| Hsa_GRHL2 | XP_011515609.1 |
| Dre_GRHL2 | NP_001076541.1 |
| Hsa_GRHL3 | NP_001181939.1 |
| Mmu_GRHL3 | XP_006538823.1 |
| Pmu_ubp | XP_038058376.1 |
| Lva_ubp | XP_041459571.1 |
| Dre_ubp | XP_005159783.1 |
| Mmu_ubp | NP_001076788.1 |
| Hsa_CP2 | XP_016859394.1 |
| FoxH tree | |
| Scl_Fox | XP_039264726.1 |
| Pzor_Fox | Polyandrocarpa_FOXH1_TRINITY_DN18220_c0_g1_i2.p1 |
| Cin_Fox | NP_001071718.1 |
| Mmu_FoxQ | NP_032265.3 |
| Hsa_FoxQ | NP_150285.3 |
| Dre_FoxQ | NP_001230273.1 |
| Apl_FoxQ | XP_022098112.1 |
| Aja_FoxQ | XP_033110155.1 |
| Scl_FoxQ | XP_039265174.1 |
| Cin_unc | XP_009861630.2 |
| Cle_Fox | Clavelina_FOXH1_TRINITY_DN386_c0_g1_i20.p1 |
| Aja_FoxF | XP_033110146.1 |
| Apl_FoxF | XP_022098116.1 |
| Mmu_FoxF | NP_034556.2 |
| Hsa_FoxF | NP_001442.2 |
| Dre_FoxF | NP_001073655.1 |
| Scl_FoxF1 | XP_039264859.1 |
| Dre_FoxH | NP_571577.1 |

| | |
|---|---|
| Hsa_FoxH | NP_003914.1 |
| Mmu_FoxH | NP_032015.1 |
| DLX tree | |
| Cin_DLX | XP_018667881.1 |
| Cle_DLX | Clavelina_DLX_TRINITY_DN3209_c0_g1_i2.p1 |
| Scl_DLX | XP_039267577.1 |
| Pzor_DLX | Polyandrocarpa_DLX_TRINITY_DN703_c0_g1_i2.p1 |
| Dre_DLX1A | NP_571380.1 |
| Mmu_DLX1 | NP_034183.1 |
| Hsa_DLX1 | NP_835221.2 |
| Dre_DLX6 | NP_571398.1 |
| Mmu_DLX6 | NP_034187.1 |
| Hsa_DLX6 | NP_005213.3 |
| Dre_DLX4a | NP_571375.1 |
| Aja_DLX6 | XP_033100592.1 |
| Apl_DLX6 | XP_022086456.1 |
| Dre_DLX5 | NP_571381.2 |
| Mmu_DLX5 | NP_034186.2 |
| Hsa_DLX5 | NP_005212.1 |
| Mmu_DLX3 | NP_034185.1 |
| Hsa_DLX3 | NP_005211.1 |
